# Engineered regulatory states resolve context-dependent genetics in macular degeneration

**DOI:** 10.64898/2026.10.09.758047

**Authors:** Eli Metzner, Karthik Guruvayurappan, Kaden M. Southard, Rico C. Ardy, Kushal K. Dey, Thomas M. Norman

**Affiliations:** Computational and Systems Biology Program, Memorial Sloan Kettering Cancer Center, New York, NY, USA; Tri-Institutional Training Program in Computational Biology & Medicine, New York, NY, USA

**Author notes:** These authors contributed equally to this work (Eli Metzner, Karthik Guruvayurappan).

## Abstract

The effects of many genes and disease-associated variants remain hidden in standard cellular models, limiting functional interpretation. Here we engineer regulatory states via CRISPR activation of transcription factors (TFs) in human RPE-1 cells, a model for age-related macular degeneration (AMD). Multiome Perturb-seq profiling of single-TF perturbations identified regulators that selectively opened chromatin at AMD GWAS variants inaccessible in resting RPE-1. Priming of partner motifs in chromatin made accessible by single-TF perturbations predicted TF pairs generating emergent transcriptional states in combinatorial Perturb-seq, providing a design principle for selecting informative interactions. Distinct TF pairs also showed variable convergence between polygenic AMD signals and Mendelian retinal disease genes, supporting an omnigenic model shaped by regulatory states. Finally, Mosaic, a chromatin-informed model of TF interaction phenotypes, nominated out-of-sample regulators, including a higher-order combination with experimentally validated emergent effects on AMD-associated programs. Engineered regulatory states thus bring hidden disease biology within experimental reach.

## INTRODUCTION

The central goal of genetics is to map genotype to phenotype, yet functional effects remain difficult to resolve despite advances in single-cell and functional genomics.^1–9^ Across 49 human tissues, 57% of GWAS associations do not colocalize with an observed cis-acting eQTL,^10^ and enhancer-gene mapping in 1,458 ENCODE biosamples left 44% without an assigned target gene.^11^ Perturbation data show the same pattern: two thirds of gene knockdowns do not produce a detectable transcriptional phenotype in genome-scale Perturb-seq,^12^ and 42% of constitutively expressed genes are never essential across the diverse cell lines in the DepMap.^13^ These gaps leave many disease associations without a molecular explanation and constrain emerging efforts to build virtual cell models,^14, 15^ which cannot learn from conditions that produce no phenotype.

At least some of this silence is conditional: in primary monocytes, 43% of genes with an eQTL showed an effect only after stimulation.^16^ This suggests an in vitro strategy for revealing latent function: apply one perturbation to move cells into a new regulatory state, then test whether a second perturbation behaves differently in that context than it does on its own. These context-dependent effects, or genetic interactions, can expose phenotypes that neither perturbation reveals individually, and combinatorial Perturb-seq provides a scalable way to measure the transcriptional phenotypes of many perturbation pairs in parallel.^17^ Informative interactions, however, are expected to be sparse within the vast space of possible combinations. A central challenge is therefore to identify, from measurements of individual perturbations, combinations that reveal new function.

Here, we focus on transcription factor (TF) interactions as a tractable subset of this problem. TFs drive large changes in cell state, and their interactions are partly mediated by binding to DNA in a sequence- and chromatin-dependent manner. Multiome Perturb-seq makes these effects observable by measuring chromatin accessibility and gene expression in each perturbed cell, capturing both the local *cis*-regulatory effects of a TF perturbation and the broader *trans*-regulatory responses that follow.^18–24^ TF perturbations are therefore well-positioned to test whether chromatin changes induced by one predict the effect of another in combination. Changes in accessibility may also reveal regulatory relationships before they are realized within the transcriptome; during development, chromatin changes anticipate later changes in gene expression.^25–28^ By analogy, if perturbing one TF opens binding sites for another, it may prime the regulatory landscape for that second TF to act in a new way. Such “chromatin priming” could identify combinations capable of producing emergent phenotypes.

The regulatory states created by TF perturbations may also expose otherwise hidden mechanisms relevant to disease genetics. Most disease-associated variants lie in noncoding regions^29^, so interpreting their effects requires eliciting conditions in which the *cis*-regulatory elements (CREs) they occupy are active. Regulatory state matters beyond these local effects, however, as it also shapes the *trans*-regulatory network through which variant effects propagate to downstream genes. Under the omnigenic model, distributed genetic effects converge through these networks on disease-proximal core genes.^2, 6, 30–36^ If network connectivity changes with regulatory state, these routes of convergence may change as well. Perturbing regulatory state may therefore inform disease genetics at two levels: in *cis*, by exposing regulatory elements harboring putative causal variants, and in *trans*, by uncovering regulatory connections to disease-proximal genes absent in cells at rest.

We test this framework in human RPE-1 cells, using age-related macular degeneration (AMD) as the primary disease model. The retinal pigment epithelium (RPE) plays a central role in AMD pathology, and major AMD risk loci implicate complement regulation and RPE function.^37, 38^ Yet many loci remain mechanistically unresolved despite increasingly detailed multiomic maps of the RPE.^39^ AMD therefore provides a useful test case: its genetics yield credible disease-associated signals but leave much of their functional basis unresolved, allowing us to ask whether experimentally induced regulatory states can advance their biological interpretation.

We perform CRISPR activation (CRISPRa) Multiome Perturb-seq against 119 TFs and combinatorial Perturb-seq against all 1,225 pairwise combinations among a subset of 50 TFs to test whether synthetic TF perturbations reveal AMD-relevant functions and regulatory relationships not apparent in the resting state. Chromatin priming measured in single perturbations enriched for TF pairs that generated novel transcriptional states when activated together, providing a principle for prioritizing informative combinations. Single perturbations also exposed regulatory elements harboring fine-mapped AMD variants that were inaccessible at baseline, while combinatorial perturbations connected common-variant AMD signal to genes implicated in Mendelian retinal disorders, consistent with their proposed role as disease-proximal core genes in the omnigenic model. Different TF combinations engaged different subsets of these genes, suggesting that the regulatory connections through which genetic effects converge on core genes depend on cell state and are incompletely captured at steady state. Finally, Mosaic, a chromatin-informed model of interaction phenotypes, nominated regulators of AMD-associated gene programs outside the pairwise screen whose effects emerged only in higher-order combinations. Together, these findings establish a strategy for revealing latent disease-relevant biology by constructing regulatory states in which it becomes observable.

## RESULTS

### A Multiome Perturb-seq CRISPRa screen quantifies TF function and cross-regulation

To deeply characterize a broad set of TFs, we performed Multiome Perturb-seq in an RPE-1 cell line stably expressing the VPH CRISPRa effector, targeting 120 TFs with 157 sgRNAs validated to produce transcriptional phenotypes in our previous work^40^ (**Fig. 1A, Table S1, Methods**). We achieved robust perturbation assignment (**Fig. S1A,** median 63.4% per GEM group) via T7-assisted ID amplification, employing an updated version of our engineered sgRNA expression vector^18^ with a GFP-KASH fusion^41^ that enables sorting of successfully transduced nuclei. After quality control (**Fig. S1B**), we recovered 25,129 single cells assigned exactly one sgRNA, detecting 119 of 120 TF perturbations and non-targeting controls. Of 157 sgRNAs in the library, 133 (targeting 104 unique genes, plus controls) were represented by at least 20 cells, despite strong positive (*PRDM16*, *MYC*) and negative (*GATA2*) fitness effects induced by perturbations (**Fig. S1C**). sgRNAs drove consistent activation of their TF targets, limited in magnitude to a physiological expression range (**Fig. 1B**). ChromVAR scoring^42^ revealed concordant motif activity across the genome (**Methods**): perturbation of TF families with known pioneering activity (e.g., FOXes) caused their cognate motifs to open, while perturbation of canonical repressors (*HIC1*, *ZEB2*) caused their motifs to close (**Fig. 1C**, diagonal). We also observed widespread cross-regulation, in which activation of a single TF affected the accessibility of the motifs of other TFs (**Fig. 1C**, off-diagonal).

**Fig. 1:**
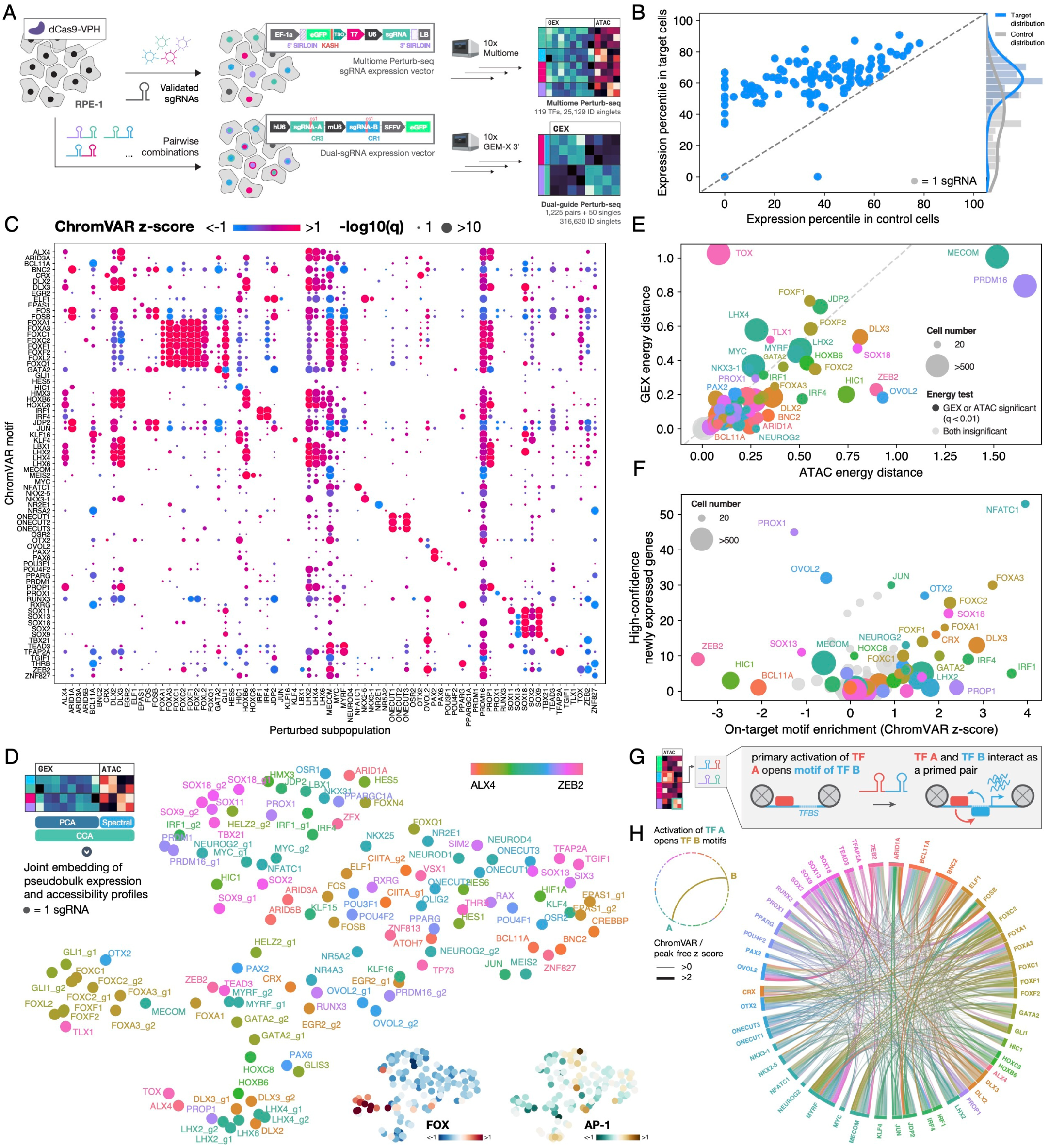
Single-TF activation with Multiome Perturb-seq organizes TF function and induces cross-regulatory chromatin phenotypes. **A.** Schematic of paired Perturb-seq experimental design with summaries of each experiment’s sgRNA expression strategy and recovered cell populations. **B.** Expression percentile of target TFs in cells containing each targeting sgRNA and all control sgRNAs. Marginal histogram shows distribution of expression percentiles for all target genes in all perturbed cells (blue) or all control cells (gray). **C.** ChromVAR motif enrichment of perturbed subpopulations (grouped by target TF) with at least one differential motif (*q <* 0.1, Kolmogorov-Smirnov test) across all motifs of target TFs. **D.** UMAP visualization of joint embedding of gene expression and chromatin phenotypes for all perturbed subpopulations, grouped by sgRNA. Points are colored by target TF in alphabetical order with a continuous color scale, such that homologs are assigned similar colors. Inset UMAPs show average activity of motifs in JASPAR FOX and AP-1 motif clusters. **E.** Energy distance calculated on normalized components of gene expression (GEX) PCA embedding and chromatin (ATAC) spectral embedding for all targets with at least 20 cells. Colored points represent targets with significant energy distance difference from control cells (*q <* 0.01, permutation test, Benjamini-Hochberg correction) in at least one modality. TFs are colored as in Fig. 1D. **F.** On-target ChromVAR motif enrichment vs. number of high-confidence newly expressed genes for all targets with at least 20 cells. High-confidence newly expressed genes are unexpressed in control cells, expressed in at least 10% of target cells, and associated with at least one concordant (differentially accessible) ATAC peak. Colored points indicate target TFs selected for dual-guide Perturb-seq screen; coloring as in Fig. 1D. **G.** Schematic of priming between illustrative TFs A and B, in which A “primes” B by opening B motifs after activation. **H.** Priming relationships among 50 TFs selected for dual-guide Perturb-seq screen. A single chord is drawn between primed TF pairs and colored by the TF whose motif is opened, with colors as in Fig. 1D. Chords between bidirectional pairs are colored by the TF whose motif has the larger z-score.

To organize the global effects of single-TF perturbations, we fit a joint embedding to gene expression and chromatin phenotypes and visualized mean populations via UMAP (**Fig. 1D, Methods**). Perturbations largely clustered by TF family, and where multiple sgRNAs against the same TF were included, they were often neighbors in embedding space, demonstrating the internal consistency of the Multiome readout. Some clusters showed specific motif activity (e.g., FOX motif accessibility in the FOX-enriched cluster), whereas motifs of broadly active TFs such as AP-1 were diffuse across the embedding.

Phenotypes driving this clustering were not equally distributed between the transcriptome and epigenome. We computed the energy distance^43^ between perturbed and control cells in each modality, normalizing embeddings so that distances could be compared directly (**Fig. 1E, Methods**). While the strengths of phenotype were largely correlated across modalities, some TFs disproportionately affected the epigenome (*ZEB2*, *OVOL2*), while others mainly affected the transcriptome (*TOX*, *LHX4*). We refined this analysis by comparing on-target motif opening in chromatin space, a proxy for pioneering activity, to the degree of *de novo* expression of genes unexpressed in control cells (**Fig. 1F**). Motif opening was neither necessary nor sufficient for *de novo* expression, suggesting that the chromatin-remodeling and downstream transcriptional effects of TFs can be partially decoupled.

Integrating these global measurements of regulatory state and function, we nominated 50 TFs for pairwise interaction profiling (**Fig. 1F** colored points, **Table S1**). Because TF binding depends on accessibility, a TF that opens the motifs of another expands the set of sites its partner can bind. We therefore hypothesized that such cross-regulatory TF pairs would tend to produce neomorphic effects on cell state, engaging regulatory programs that neither TF activates alone. We called this relationship “chromatin priming,” and the participating TFs a “primed pair,” drawing analogy to sequential priming of regulatory elements previously described in developmental biology^25–28^ (**Fig. 1G**). To comprehensively annotate primed pairs in our Multiome data, we complemented peak-based ChromVAR motif scores with a peak-free single-cell scoring pipeline (**Table S2, Methods**), identifying 539 primed pairs among these 50 TFs (44.0% of 1,225 pairs) (**Fig. 1H, S1D, S1E**). Many TFs primed motifs of multiple TF families, and 57 of 539 primed pairs were “bidirectional,” where activation of each TF in a pair caused opening of its partner’s motifs.

### TF cross-regulation underlies neomorphic genetic interactions

We cloned a dual-sgRNA library targeting the nominated TF pairs, single perturbations (targeting sgRNA plus control sgRNA), and non-targeting control cells (control sgRNA plus control sgRNA) and performed direct-capture Perturb-seq in RPE-1 VPH cells (**Fig. 1A, Methods**). After quality control and ID assignment, we recovered 316,630 cells for downstream analysis (**Fig. S2A, S2B, Methods**). We observed robust activation of TFs targeted by sgRNAs in both positions of the expression vector, and activation was highly correlated between positions across TFs (Pearson *r* = 0.81), despite a trend toward weaker activation in the 3’ “B” position (**Fig. 2A**). Consequently, pairs targeting the same TFs in opposite orientation yielded highly correlated transcriptional profiles compared to random pairs targeting different genes (**Fig. S2C,** median *r* in PCA space = 0.58 vs. 0.03). We therefore collapsed both orientations into a composite single phenotype for each pair. In the final cell population, all pairs were represented by at least 12 cells (median 171), all single perturbations by at least 77 cells (median 475), and non-targeting controls by 4,566 cells (**Fig. S2D**).

**Fig. 2:**
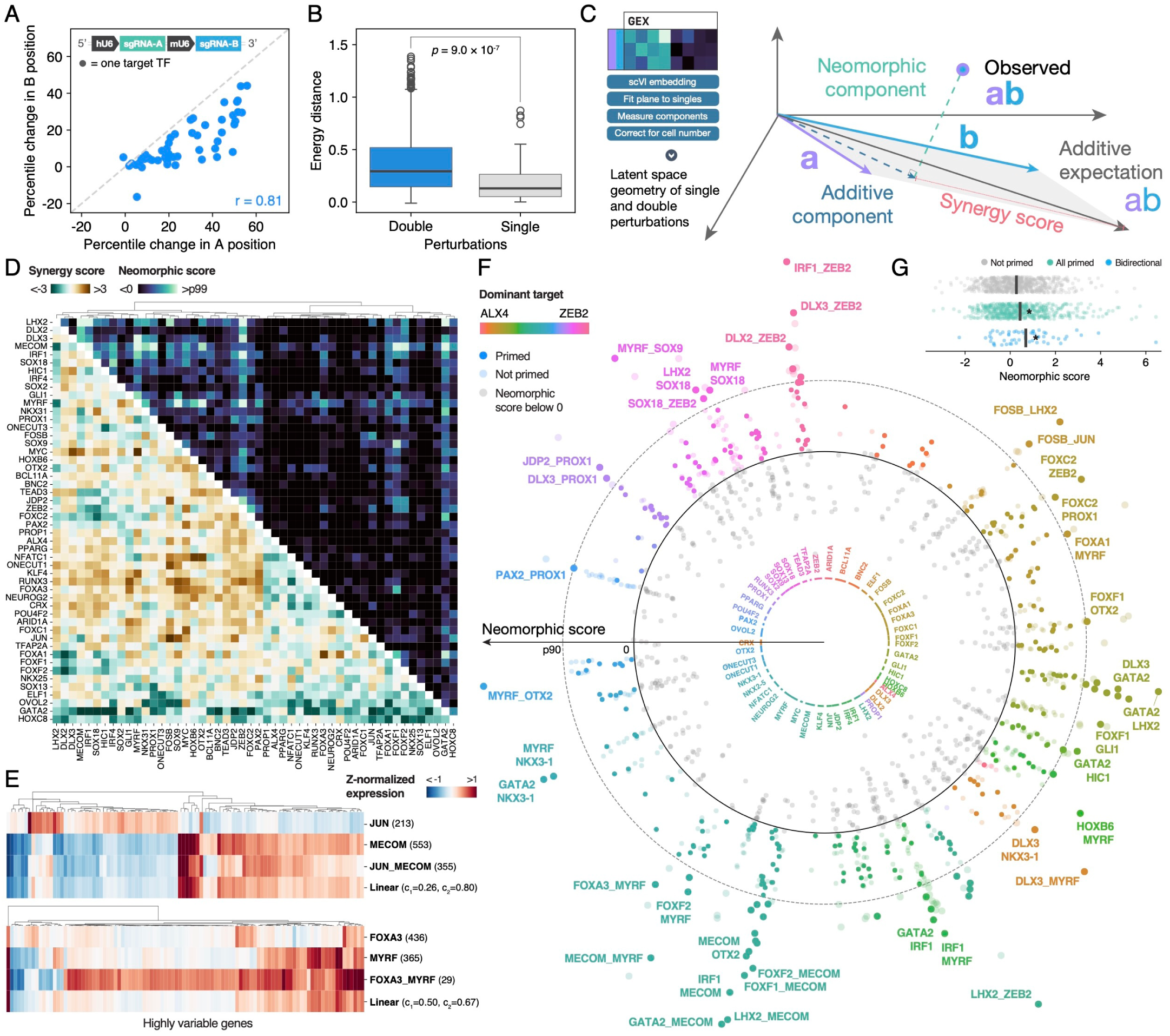
Combinatorial TF activation connects neomorphic TF pairs to chromatin states observable in single-TF perturbations. **A.** Expression percentile change for targeting sgRNAs in each position of the dual-sgRNA expression vector (depicted at upper left). Percentile change calculated over all cells containing each target TF’s sgRNA in the indicated position, relative to control cells containing NTC sgRNAs in both positions. **B.** Energy distance to control cells calculated in normalized PCA space for all double and single perturbations. *P* -value via one-sided Mann-Whitney test. **C.** Schematic of embedding strategy to quantify nonlinearity of TF interactions via latent space geometry. **D.** Genetic interaction maps quantifying pairwise TF phenotypes via synergy score (below diagonal) and neomorphic score (above diagonal). TFs are clustered by synergy score using cosine distance and Ward’s linkage. **E.** Z-normalized expression of highly variable genes for single perturbations and double perturbation, compared to prediction of linear model fit to single perturbation expression, for pairs JUN_MECOM and FOXA3_MYRF. **F.** Neomorphic scores of all TF pairs grouped by dominant target. TF coloring for pairs with neomorphic score greater than 0 (solid circle) as in Fig. 1. Pairs with neomorphic score above the 90^th^ percentile across all pairs (dashed circle) appear as larger points. Width of each TF’s sector, as in Fig. 1H, is proportional to number of priming connections. **G.** Neomorphic scores of 686 non-primed, 539 primed, and 57 bidirectional pairs. Large tick mark denotes the mean neomorphic score in each group. Bidirectional pairs are a subset of all primed pairs. \**p <* 0.05, one-sided Welch’s t-test.

To quantify the diversity of transcriptional states reached by perturbing TFs in combination, we compared the transcriptional profiles of double and single perturbations using energy distance (**Methods**). Double perturbations were significantly further from control cells than single perturbations in this metric (**Fig. 2B**, *p* = 9.0 × 10^-7^). To assess the biological relevance of this result, we compared perturbation phenotypes to five in vivo RPE cell states from a recent single-cell atlas of human RPE and choroid^44^ (**Fig. S3A**) by scoring perturbed subpopulations for expression of marker genes uniquely upregulated in each state (**Methods**). Across all RPE states, double perturbations induced more variance in state scores than single perturbations (**Fig. S3B,** *q <* 0.05). We explored this further in pairs involving the RPE identity factor *MYRF*.^45^ Combinatorial activation of *MYRF* and *PROX1* uniquely induced marker genes of the *CST3*+ senescence-associated RPE state that were not induced via MYRF perturbation alone, while perturbation of *MYRF* and *FOXA1*, and *MYRF* and *CRX*, respectively, induced markers of the *INSYN2B*+ and *LGI1*+ states, whose accessible chromatin in the atlas was specifically enriched for AMD risk loci (**Fig. S3C**). Combinatorial TF activation therefore generates stronger effects on cell state than single-TF perturbations and can access states observed in vivo that single-TF perturbations cannot.

To characterize modes of genetic interaction among TF pairs, we embedded transcriptional profiles with scVI^46^ and used the geometry of this latent space to quantify interaction linearity and synergy (**Fig. 2C, Table S3, Methods**). We resolved the coordinates of each double perturbation AB into an additive component within the plane defined by single perturbations A and B and an orthogonal component capturing transcriptional behavior present in neither single. We defined a *neomorphic score* from the orthogonal component to quantify emergence of a new transcriptional direction upon double perturbation, and a *synergy score* from the additive component to quantify whether the magnitude of the double phenotype was stronger or weaker than expected. We also assigned the single TF A or B with greater cosine similarity to AB as AB’s *dominant target*. Visualizing combinatorial phenotypes as genetic interaction maps of neomorphic and synergy scores (**Fig. 2D**), hierarchical clustering revealed shared structure between the two, indicating that linear and nonlinear deviations from pure additivity often co-occur, though we also observed unique behavior in both metrics. As in previous work,^17^ we fit linear models to decompose double transcriptional profiles into effects attributed to each single (**Methods**). Model coefficients often agreed with our embedding-based scores, and we could find clear examples nominated by both strategies as, e.g., buffering (*JUN_MECOM*, synergy score -1.76) or neomorphic (*FOXA3_MYRF*, neomorphic score 2.90) (**Fig. 2E**).

Grouping pairs by dominant target, we observed that several TFs with strong effects on chromatin state alone (**Fig. 1E**) induced highly neomorphic interactions across many partners (e.g., *MECOM*, *ZEB2*) (**Fig. 2F**). Primed TF pairs were enriched in the upper tail of the neomorphic score distribution (enrichment in top 10%: OR = 1.48, *p* = 0.024), and had higher neomorphic scores overall (**Fig. 2G**, mean score 0.44 vs. 0.27, *p* = 0.012), an effect that was more pronounced for bidirectional pairs (mean score 0.69, *p* = 0.017). Primed TF pairs therefore interact more neomorphically when activated together, indicating that motif-level cross-regulation anticipates emergent transcriptional states. Critically, primed TF pairs can be identified from single-TF phenotypes alone, scaling linearly rather than quadratically with the number of TFs considered. Priming therefore provides a simple yet powerful design principle to enrich for TF combinations that produce novel transcriptional outcomes, a problem for which few prospective criteria currently exist.

### TF activation reveals cryptic regulatory elements at AMD risk loci

The priming result can be viewed as one instance of a broadly useful property of TF perturbations: they are a simple way to impose defined and programmable regulatory states that modify how other genetic effects manifest. We reasoned that this could be exploited to study disease-associated variants, which are mostly noncoding^29, 47–49^ and therefore inert unless the elements they occupy are active – a condition that may not be met in cultured cells in vitro. We therefore asked whether TF activation could expose regulatory elements harboring AMD risk variants that are inaccessible in resting cells.

To assess the relevance of RPE-1 for AMD genetics, we first tested whether fine-mapped AMD risk variants preferentially overlapped accessible chromatin in RPE. Of 301 fine-mapped (PIP *>* 0.1) variants from a recent European-ancestry AMD meta-analysis^50^ (*n* = 57,290 cases, 324,430 controls), overlap was greatest with RPE peaks (enrichment = 5.29-fold, recall = 0.090) relative to other choroidal cell types (enrichment = 2.50-3.97-fold, recall = 0.013-0.060), consistent with the central role of RPE in AMD (**Fig. S4A**).^39^ This cell-type-specificity was further supported by heritability enrichment analysis (**Fig. S4B, Methods**).^48^ Bulk ATAC-seq peaks in RPE-1^40^ also showed significant enrichment for AMD heritability (**Fig. S4C**; enrichment = 6.93-fold, *p* = 0.018). Together, these analyses establish RPE as a relevant cell type and RPE-1 as a suitable experimental model for studying AMD genetics. Even so, RPE and RPE-1 peaks covered only 9.0% and 5.3% of all fine-mapped variants, respectively, leaving most AMD risk variants outside accessible chromatin in resting RPE-1.

We defined perturbation-specific accessible regions as peaks present in one or more TF activation contexts but not detected in control cells (**Fig. 3A, Methods**). These regions recovered 28% of AMD GWAS fine-mapped variants, compared with 5 to 7% recovered by peaks from control cells or bulk RPE-1 ATAC-seq^51, 52^, while showing comparable enrichment relative to a background of common and low-frequency variants^53^ (**Fig. 3B**, enrichment = 1.63-fold, *p* = 2.2 × 10^-7^ vs. 1.58- and 1.61-fold). Restricting perturbation-specific regions to elements linked to candidate target genes by scE2G^54^ (**Fig. 3A, Methods**) increased enrichment to 3.67-fold (vs. 2.42-fold in control cells) at the cost of recall, consistent with peak–gene linking selecting for elements with greater regulatory consequence.^54, 55^ Overall, 75 fine-mapped AMD variants (PIP *>* 0.1) overlapped perturbation-specific regions but no control or bulk RPE-1 peak, including 23 variants linked by scE2G to at least one candidate target gene (**Fig. 3C, Fig. S5A**). For example, the AMD risk variants rs376659441 and rs113616316 (both PIP = 1) lie within an intronic *ME3* peak that becomes accessible following *PAX2* activation and was linked to *ME3* by scE2G (**Fig. 3D**). TF activation with chromatin accessibility readout therefore provides a scalable strategy for revealing accessible regions at otherwise quiescent AMD disease risk variants.

**Fig. 3:**
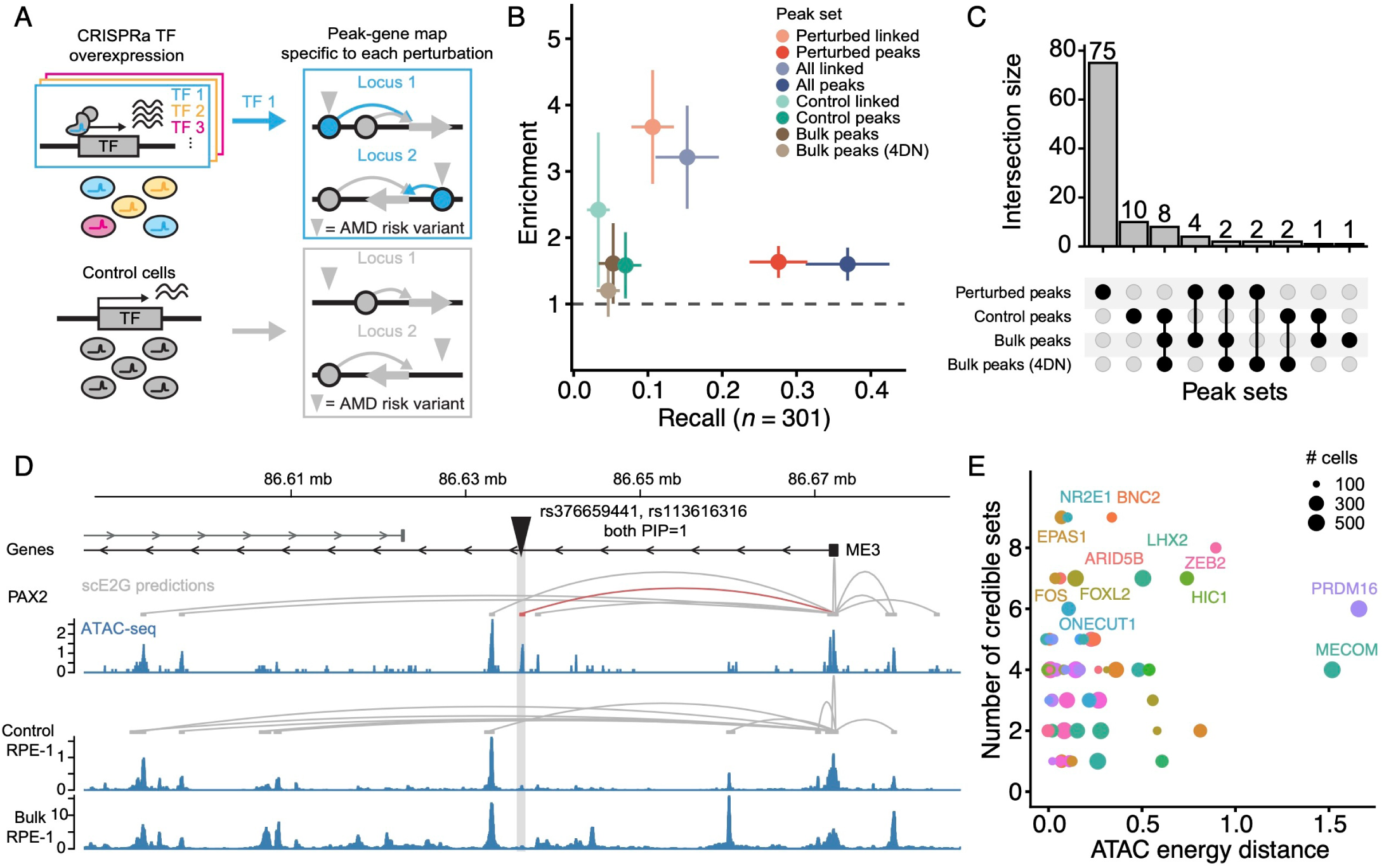
Single-TF activation with Multiome Perturb-seq reveals perturbation-induced accessible chromatin overlapping AMD risk loci. **A.** Schematic of procedure to generate peak sets, peak-gene maps, and assess their overlap with AMD risk variants for each perturbation in the Multiome Perturb-seq experiment. **B.** Enrichment vs. recall of 301 fine-mapped AMD risk variants (PIP *>* 0.1) by different peak sets derived from RPE-1 cells. “Linked” sets - peaks linked to genes using scE2G. “Perturbed” sets - union of peaks across all perturbations, removing overlaps with control peaks. “All” sets - union of all peaks across all perturbations and control cells. Dots - mean. Error bars - 1.6 * s.e. **C.** UpSet plot with recall of AMD fine-mapped variants by different peak sets. Peak sets defined as in Fig. 3B. **D.** Browser plot of the *ME3* locus. Tracks (top to bottom): genes, peak-gene links for *PAX2* CRISPRa cells, ATAC-seq signal for *PAX2* CRISPRa cells, peak-gene links for control cells, ATAC-seq signal for control cells, and ATAC-seq signal for bulk RPE-1 cells from 4DN. **E.** Number of perturbation-specific credible sets revealed vs. ATAC energy distance of perturbations with *>* 100 cells (*n* = 54) from the Multiome Perturb-seq experiment.

We next assessed whether this recovery of AMD risk variants was driven by a small number of perturbations or distributed across diverse TF-induced regulatory contexts. We defined perturbation-specific credible sets as those in which at least one variant (PIP *>* 0.1) overlapped a perturbation-specific region and none overlapped regions accessible in control cells (**Methods**). All 54 TF targets with at least 100 cells revealed at least one perturbation-specific credible set (**Fig. 3E**). Among AMD variants recovered within the 62 perturbation-specific credible sets, 45% were exposed by a single TF perturbation and 29% by five or more (**Fig. S5B**). The number of credible sets revealed per perturbation correlated minimally with the number of cells profiled (*r* = 0.05) or with energy distance in ATAC embedding space (*r* = 0.12), indicating that recovery of disease variants reflects perturbation-specific responses rather than simply the magnitude of global chromatin remodeling (**Fig. 3E**).

Finally, because individual TF perturbations sometimes co-activated regulatory elements harboring variants from distinct AMD fine-mapped loci, we explored targeted genetic interaction testing at these locus pairs. This analysis identified suggestive genotype-by-genotype associations, which we consider hypothesis-generating (**Fig. S6, Methods**). Together, these results show that TFs chosen without regard to AMD nevertheless impose altered regulatory contexts that can expose many risk variants silent at baseline. TF Multiome Perturb-seq may therefore offer a general strategy for interpreting noncoding disease variation even when precise regulatory drivers are unknown.

### Combinatorial TF perturbations reveal state-dependent convergence of polygenic AMD and Mendelian retinal disease genes

Having shown that single-TF perturbations expose AMD risk variants in cis, we next asked whether the paired Perturb-seq data could reveal how variant effects propagate in trans through their downstream genes. Because combinatorial perturbations drove stronger and more diverse changes in cell state than single perturbations (**Fig. 2B, Fig. S3B**), we reasoned that they might also expose regulatory connections to AMD genes that neither single perturbation reveals. We therefore sought to assess whether AMD disease genes were more likely to be regulated non-additively by the TF pairs in our dataset. For each TF pair, we quantified each gene’s interaction phenotype as its residual from an additive expectation derived from its two single-TF perturbations, accounting for measurement uncertainty across conditions (**Fig. 4A, Methods**). We defined the *Pooled Combinatorial Interaction (PCI) score* of each gene as the L_2_ norm of its residuals across all 1,225 double perturbations, providing a global measure of sensitivity to non-additive combinatorial regulation (**Fig. 4A, Methods**).

**Fig. 4:**
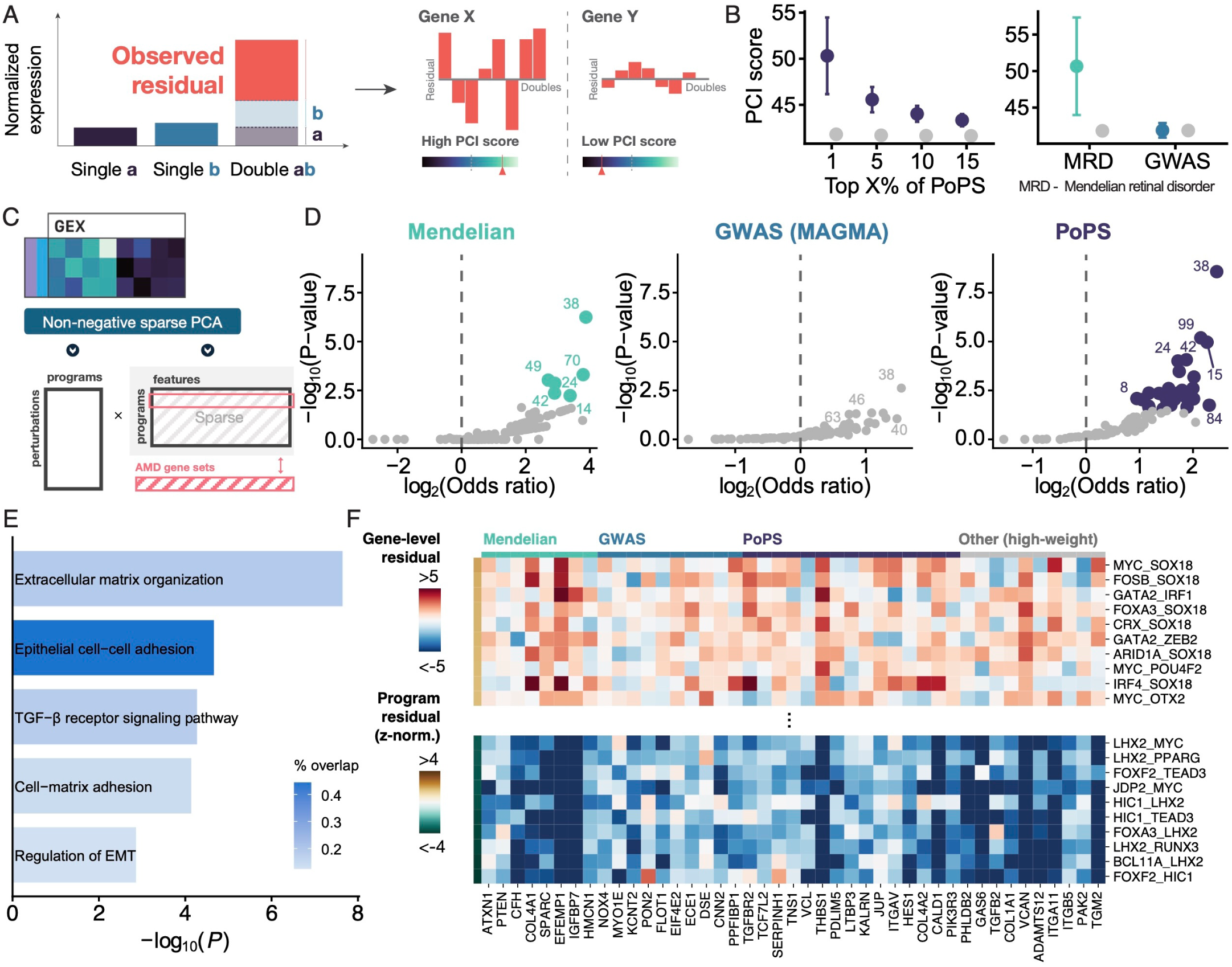
Combinatorial TF CRISPRa Perturb-seq reveals enrichment of MRD and polygenic AMD disease genes among perturbation-responsive genes and programs. **A.** Schematic of residual quantification for each double TF perturbation. **B.** PCI scores of different disease gene sets vs. genes outside the disease gene set. Colored points - mean for gene set genes. Gray points - mean for all other genes. Error bars - 1.96 * s.e. **C.** Schematic of gene program creation using non-negative sparse PCA. **D.** Enrichment of gene programs for AMD disease gene sets. *P* -values computed using a Fisher’s exact test. **E.** Pathway enrichments for program 38 genes against all program genes using Enrichr^74–76^ with GO Biological Process 2026.^77, 78^ % overlap - proportion of pathway genes in program 38. *P* -values are from a Fisher’s exact test. **F.** Residuals of MRD genes, AMD GWAS genes, AMD PoPS genes, and other high-weight genes for program 38. Rows show doubles with top and bottom 10 program-level residuals.

We then compared PCI scores of AMD genes to those of matched background genes, using three complementary gene sets: GWAS genes (prioritized using MAGMA^56^); Polygenic Priority Score^57^ (PoPS) genes demonstrating functional convergence with GWAS genes; and Mendelian retinal disorder (MRD) genes for which AMD is a phenotypically matched complex disease^58^ (**Table S4, Methods**). We observed low to moderate overlap across these AMD gene sets (Jaccard index 0.0054-0.073). MRD genes and AMD PoPS genes exhibited substantially greater PCI scores than matched background genes (**Fig. 4B**, 50.7 versus 41.8, *p* = 7.0 × 10^-3^ for monogenic, 46.8 vs. 41.7, *p* = 4.2 × 10^-8^ for top 500 PoPS genes), whereas MAGMA-prioritized genes showed no increase (41.9 versus 41.9, *p* = 0.48). This pattern was broadly distributed across TF pairs: 37% of pairs showed significantly (nominal *p <* 0.05) elevated interaction residuals among the top 500 PoPS genes and 23% among MRD genes, compared with only 3% among MAGMA-prioritized genes. PoPS genes for AMD also had the 4^th^-largest elevation of PCI scores when compared to 113 other complex traits (**Fig. S7, Methods**). MRD and PoPS AMD genes are therefore disproportionately sensitive to combinatorial TF regulation, whereas genes nominated by GWAS proximity alone are not.

We next asked whether MRD and PoPS AMD genes, despite overlapping little as sets, might be coupled through shared transcriptional programs. We factored the expression matrix via non-negative sparse PCA^40^, producing 100 transcriptional programs defined by sparse co-regulated gene features weighted by importance (**Fig. 4C, Table S5, Methods**). Six programs were significantly enriched for MRD genes and 30 for PoPS-prioritized genes, whereas none were enriched for MAGMA-prioritized genes (*q <* 0.1, **Fig. 4D**). Five of the six MRD-enriched programs were also enriched for PoPS AMD genes, indicating pathway-level convergence between monogenic and polygenic disease genes. Program 38 showed the strongest odds ratio for both MRD and top 500 PoPS AMD genes (OR = 14.69, *p* = 5.7 × 10^-7^ and OR = 5.44, *p* = 2.7 × 10^-9^ respectively); it also ranked first in odds ratio for MAGMA-prioritized genes, although this association did not reach significance after multiple testing correction (OR = 2.92, *p* = 2.4 × 10^-3^). Genes in program 38 showed enrichment in extracellular matrix organization, epithelial cell-cell adhesion, and regulation of epithelial-to-mesenchymal transition (**Fig. 4E, Methods**). TF pairs showed opposing effects on program 38, with SOX18-containing combinations prominent among positive regulators and LHX2- and HIC1-containing combinations among negative regulators (**Fig. 4F**). Program 38 thus marks a perturbation-responsive gene program in which MRD and polygenic AMD genetic signals converge, which we refer to hereafter as the AMD convergence program.

Conventional PoPS relies on gene features measured largely at baseline, such as expression atlases, pathways, and protein interactions. Because AMD genes were highly responsive to our combinatorial perturbations, we reasoned that perturbation responses in a disease-relevant cell type might better connect common-variant association to disease genes. We therefore adapted the PoPS framework to use features from our Perturb-seq experiment (**Fig. 5A, Methods**), referring to these adapted PoPS scores as “Perturb-PoPS” scores. Because monogenic disease status was not used in training, recovery of MRD genes could serve as an independent evaluation of this approach. Despite deriving from a single cellular system and using substantially fewer features than conventional PoPS, Perturb-PoPS scores showed 3.38-fold enrichment in MRD genes (*p* = 1.0 × 10^-3^) among the top 500 predictions, exceeding the original PoPS model (**Fig. 5B**). Perturb-PoPS recovered 13 of 41 MRD genes among its top 1000 predictions, compared with 8 using conventional PoPS or a PoPS model trained on features derived from control RPE-1 cells (**Fig. S8A**). Of the 13 MRD genes recovered among the top 1,000 Perturb-PoPS predictions, 5 (38%) were not among the top 1,000 genes using either conventional PoPS or control-RPE1-derived features. These re-prioritized MRD genes include *MMP2* (PoPS rank: 7,433, Perturb-PoPS rank: 29), *HMCN1* (PoPS rank: 6,523, Perturb-PoPS rank: 613), and *RAB31* (PoPS rank: 5,317, Perturb-PoPS rank: 819), none of which has direct GWAS support based on MAGMA. Notably, this convergence was stronger when including features from TF pairs, suggesting that combinatorial TF activation provides disease signal beyond the extent of single-TF perturbation (**Fig. S8B**). It was not observed when MAGMA scores were shuffled, suggesting that it does not simply reflect co-regulation of monogenic genes with any perturbation-responsive gene (**Fig. S8C**). Many genes prioritized by Perturb-PoPS were also members of the AMD convergence program, so unsupervised program discovery and GWAS-guided prediction both independently point to its importance.

**Fig. 5:**
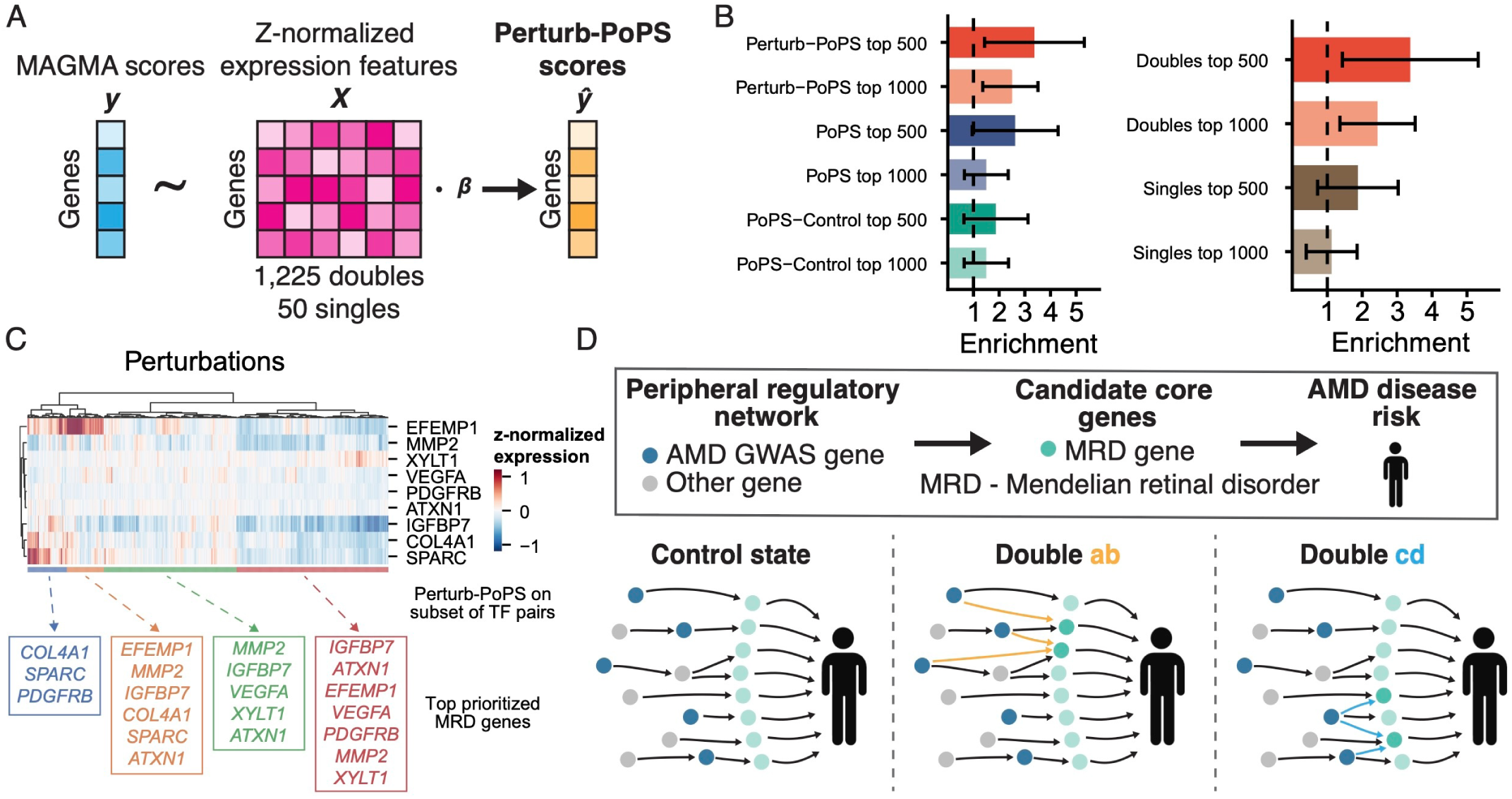
Convergence of polygenic AMD and Mendelian retinal disease genes in dual guide Perturb-seq is state-specific. **A.** Schematic for computing the Perturb-PoPS score. **B.** Enrichment of MRD genes by different PoPS score variations. Perturb-PoPS - training features from our Perturb-seq data. PoPS-control - training features from control RPE-1 cells. PoPS - training features from original PoPS feature set. Enrichments shown for either top 500 or top 1000 genes per score. Doubles - training features from singles and doubles. Singles - training features from only singles. Error bars - 1.6 * s.e. **C.** Gene expression for MRD genes with a top 500 Perturb-PoPS score. Boxes below display MRD genes in the top 1000 Perturb-PoPS genes when running Perturb-PoPS exclusively on TF pairs within that cluster. **D.** Schematic of perturbation state-specific AMD polygenic and Mendelian retinal disorder gene convergence.

Mendelian disease genes have been proposed as empirical proxies for the core genes of the omnigenic model, and this idea has been tested empirically in blood traits, where Mendelian disease genes showed properties expected of core genes in steady-state gene coexpression and regulatory networks.^32, 33^ MRD genes display two analogous properties in our screen: they are unusually sensitive to non-additive regulation across diverse TF combinations (**Fig. 4B**), and common-variant AMD association signals preferentially converge on them when propagated through perturbation-derived regulatory features (**Fig. 5B**). Importantly, this convergence was weaker when assessed using features derived from control cells, indicating that it is not fully captured by steady-state regulatory information.

Since our paired perturbation screen tests each TF perturbation with 49 partners, it uniquely allows us to assess whether the regulatory coupling of polygenic AMD genes and disease-proximal MRD genes is fixed across cellular states. We focused on 9 MRD genes strongly prioritized by Perturb-PoPS (among top 500 genes) and quantified their responses across all 1,225 TF pairs (**Fig. 5C**). Different TF combinations affected markedly different subsets of these genes. Even the most broadly engaged genes, *EFEMP1*, *IGFBP7*, and *SPARC*, varied substantially in response direction and magnitude across state. Running Perturb-PoPS on different subsets of TF-TF pairs prioritized different subsets of these 9 candidate core genes (**Fig. 5C**). Together, these observations reveal two levels of state dependence in the regulatory architecture linking polygenic and monogenic disease genetics (**Fig. 5D**). First, combinatorial TF perturbations can expose polygenic-to-candidate-core convergence that is not apparent from steady-state regulatory networks. Second, the routes of convergence can differ substantially across these TF perturbations. Thus, steady-state networks provide only a limited snapshot of the routes through which polygenic effects can converge on disease-proximal genes, while TF perturbations can reveal alternative regulatory paths that together define a state-dependent architecture of omnigenic model convergence (**Supplemental Note**).

### Mosaic predicts higher-order TF combinations that modulate AMD gene programs

The preceding analyses identify TF combinations that engage AMD-associated programs, but do not explain why they do so. We reasoned that TF interactions might be unusually interpretable: they are mediated by binding to chromatin, which the Multiome readout measures directly, and the priming result indicates that single-TF chromatin phenotypes can anticipate them in some cases. We therefore developed a model, Mosaic (**Mo**tif **S**ignatures of **A**ccessibility-**I**nduced **C**ooperativity), to test whether these phenotypes could predict interaction effects at individual genes, which would both explain how interactions arise and potentially nominate higher-order combinations not profiled in the screen. Mosaic predicts each gene’s interaction residual for a TF pair from the *cis*- and *trans*-chromatin environments induced by each constituent TF (**Fig. 6A, Methods**). Here, a gene’s *cis*-chromatin environment comprises the ATAC-seq peaks near its TSS described by TF motif, CRE identity, and single-perturbation accessibility features, as well as the gene’s expression changes following single-TF perturbations. A TF’s *trans*-chromatin environment comprises the global changes in motif accessibility induced via its activation.

**Fig. 6:**
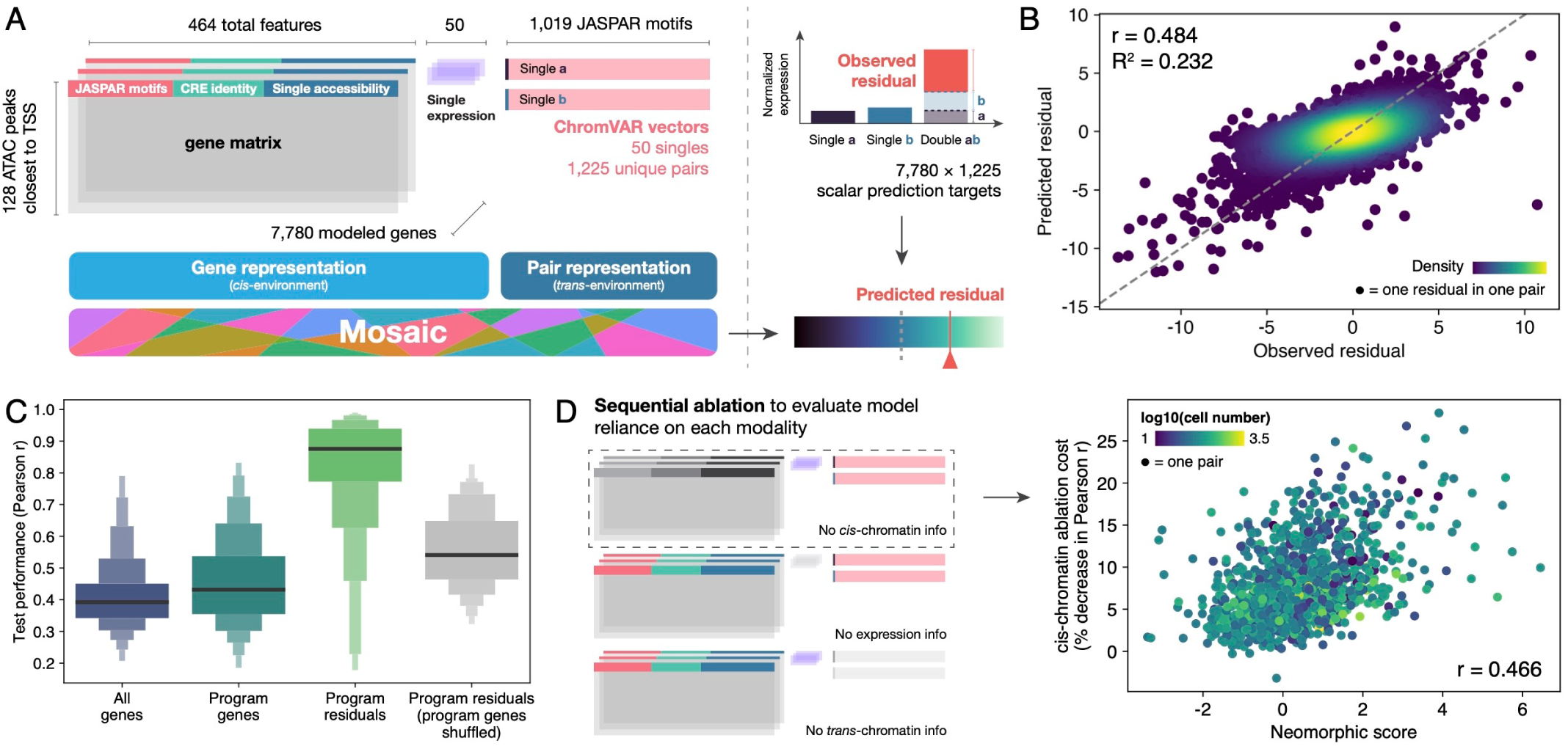
Mosaic predicts interaction phenotypes from *cis* and *trans-*chromatin states induced by single-TF perturbations. **A.** Schematic of interaction modeling framework, inputs, and outputs. **B.** Observed vs. predicted interaction residuals for 1,225 TF pairs across held-out genes in 12 cross-validation folds. 1% of predictions were sampled for plotting. Correlation and R^2^ values reflect predictions across all held-out genes. **C.** Distributions of Pearson correlation of observed and predicted residuals across all pairs for all held-out genes and program genes; program-level residuals for all programs; and program-level residuals calculated with each gene’s program label randomly permuted. **D.** Schematic of ablation strategy to test model reliance on each input modality (left); neomorphic score vs. *cis*-chromatin ablation cost (percent decrease in Pearson correlation of observed vs. predicted residuals across all held-out genes when ablating cis-chromatin information) for each TF pair (right).

Built on a Transformer architecture^59, 60^, Mosaic learns which sequence features near each gene matter for a given TF pair, weighting them according to the chromatin changes induced by the pair’s single TFs (**Fig. S9A, Methods**). It outperformed simpler bilinear and neural network models (**Fig. S9B, S9C**), and in chromosome-level cross-validation, predicted interaction residuals of held-out genes with a Pearson correlation of 0.484 (**Fig. 6B**). As in our previous work^18^, program genes, which parametrize the major axes of perturbation-induced variation in the experiment, were modeled with higher accuracy than non-program genes (**Fig. 6C**). We modeled residuals at program level by aggregating predicted and observed gene-level residuals using each gene’s program weights. Though this “de-noising” procedure increases performance even for random subsets of genes, performance across programs was substantially higher than across randomly selected genes (**Fig. 6C**, median *r* = 0.878 vs. 0.515), suggesting that shared regulation between co-varying sets of program genes provides a coherent and noise-robust modeling substrate. To assess the unique information content of each modality, we performed sequential ablation and measured changes in performance (**Fig. 6D**). Ablating single-perturbation expression or *trans*-chromatin information sharply degraded performance globally (**Fig. S9D**). By contrast, the cost of ablating *cis*-chromatin features varied by pair and increased with its neomorphic score (Pearson *r* = 0.466, *p* = 5.7 × 10^-67^). Local regulatory context is thus disproportionately informative for predicting the most nonlinear genetic interactions.

To nominate regulators of gene programs, we computed attributions for Mosaic’s residual predictions, which score how much each chromatin feature in nearby peaks contributes to the predicted residual for each gene and pair (**Fig. 7A, Methods**). Aggregating these scores across program genes, we distinguished constitutive regulators, whose attribution remains high across many TF-pair contexts, from conditional regulators, whose importance depends on the specific upstream TF combination. Hierarchical clustering of attribution scores for the AMD convergence program grouped pairs by their observed program-level residuals, implying that many chromatin features differ in importance between pairs that induce or repress the program (**Fig. 7B**). Mosaic nominated TF families associated with the ER stress response as top constitutive regulators of the program, including EGR, CREB3L3, NFKB, CREB1, and HSF. The ER stress response has been implicated in AMD, and direct evidence links ER stress and chronic activation of the NFKB pathway to retinal pathology.^61–63^ Notably, motifs of CREB3L3, a CREB-family TF canonically involved in ER stress, inflammation, and lipid metabolism,^64, 65^ were highlighted in a recent Multiome atlas of the human retina as differentially accessible in choroid from aging vs. young donors, consistent with a role in the age-dependent retinal pathology characteristic of AMD.^44^ Negative constitutive regulators included homeodomain, RUNX, and CUX-family motifs, as well as sgMYRF accessibility and motifs of SREBF1, a TF known to functionally antagonize CREB3L3.^66^ Conditional regulators aligned with TFs observed to induce the program (**Fig. 4F**), including SOX, IRF, and MYC motifs, sgGATA2 accessibility, and sgSOX18 accessibility. Other notable conditional regulators included motifs of AP-1, also recently implicated in AMD pathology^44^, and BARX2, a homeobox TF known to regulate extracellular matrix remodeling and cell adhesion in eye development^67^ (**Fig. 7C**).

**Fig. 7:**
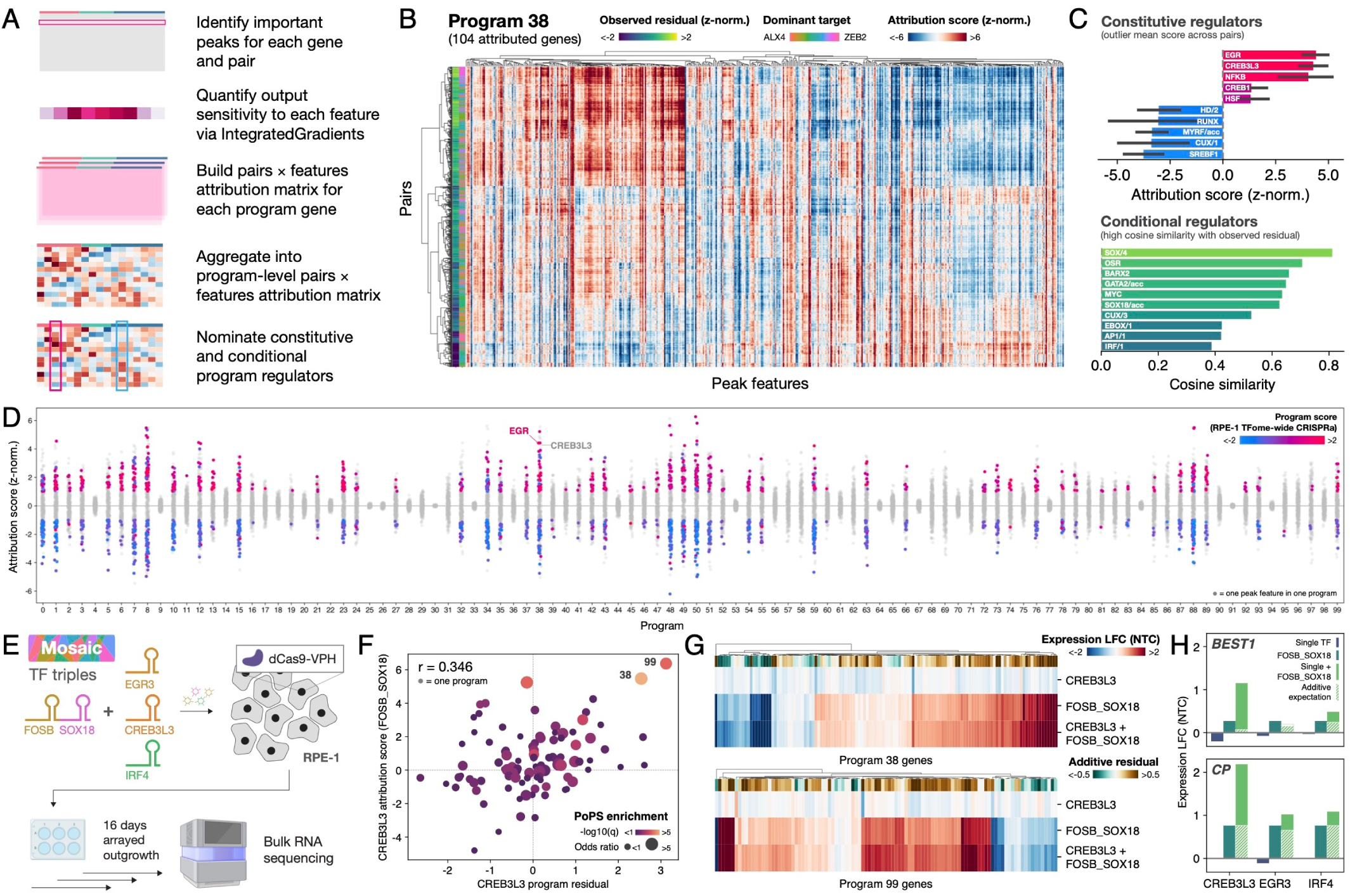
Attribution of Mosaic predictions identifies higher-order TF combinations that modulate AMD gene programs. **A.** Schematic of feature attribution strategy to nominate TF regulators of each sparse expression program **B.** Pairs-by-features attribution matrix for Program 38. Rows and columns are hierarchically clustered using correlation distance and Ward’s linkage. Dominant target labels as in Fig. 2F. **C.** (Top) mean attribution score across all pairs of select constitutive regulators of Program 38; error bars show IQR. (Bottom) cosine similarity of attribution score vector with observed program-level residuals for select conditional regulators. **D.** Mean attribution scores across pairs for all features in each program. Points are colored by program score of populations in our previous TFome-wide RPE-1 CRISPRa screen assigned to each feature. Features where no population has a program score with absolute value greater than 1, or where absolute value of mean attribution score was less than 1, are colored grey. **E.** Schematic of arrayed screen to measure transcriptional states resulting from triple-TF perturbations nominated by the model. **F.** Program attribution score for CREB3L3 motifs with respect to FOSB_SOX18 predictions vs. program residual in the CREB3L3 triple-TF condition. Programs are labeled with odds ratios and q-values of overlap with AMD PoPS genes as in Fig. 4D. **G.** Log fold changes in gene expression vs. NTC (Expression LFC) in CREB3L3 single, FOSB_SOX18 double, and triple-TF conditions across all genes in Programs 38 and 99. **H.** Expression LFC for MRD genes *BEST1* and *CP* in single-only, double-only, and triple-TF conditions. Hatched bars show additive expectation (double + single). *CP* expression was undetected in the *IRF4* single condition, so its additive expectation reflects only double expression.

We next asked whether these attributions recovered regulators in an orthogonal dataset, reasoning that that attribution scores for a program’s constitutive regulators would align with effects of activating those TFs individually in our previous TFome-wide RPE-1 CRISPRa screen.^40^ In 63 of 100 programs, at least one TF’s CRISPRa effect was concordant with Mosaic’s attribution score for its motif or accessibility (**Fig. 7D, Methods**). Notably, while HSF, EGR, and NFKB TFs all induced concordant effects on the AMD convergence program when activated individually, CREB3L3 did not, suggesting that its effect on the program might only emerge in the context of a higher-order TF interaction. Encouraged by this result, we used Mosaic attributions to rationally design triple-TF perturbations. Each combined the double FOSB_SOX18, which upregulated the AMD convergence program with positive interaction residuals for many high-weight program genes (**Fig. 4F**), with one of three TFs. EGR3 and CREB3L3 were not perturbed in the dual-sgRNA screen and were nominated solely through their motifs; IRF4 synergized with SOX18 across Program 38 genes (**Fig. 4F**) but had minimal effects as a single regulator. We implemented these TF triples in arrayed format and profiled each condition by bulk RNA sequencing (**Fig. 7E, Methods**).

We quantified non-additive interactions between FOSB_SOX18 and each single TF by calculating residuals at gene and program level (**Methods**). Consistent with Mosaic’s predictions, the AMD convergence program had a positive program residual in all three conditions, and across all programs, residuals in the CREB3L3 triple-TF condition correlated with attribution scores for CREB3L3 motifs in the FOSB_SOX18 background (Pearson *r* = 0.346, *p* = 4.2 × 10^-4^), although this relationship was weaker for EGR3 and IRF4 (**Fig. 7F, S10A, S10B**). In the convergence program and the similarly AMD-associated Program 99, CREB3L3, whose single phenotype was largely silent, dramatically enhanced the effect of FOSB_SOX18, inducing larger expression changes in the triple-TF condition than predicted by an additive expectation of double and single conditions (**Fig. 7G**). Strikingly, the CREB3L3 triple-TF condition uniquely upregulated MRD genes *BEST1* and *CP*, which were detected in too few single cells in the Perturb-seq screen for modeling or program creation (**Fig. 7H**). Both genes ranked within the top 0.4% of residuals among 11,255 protein-coding genes detected in single, double, and triple conditions (**Fig. S10C**). Together, these results position CREB3L3 as a regulator of AMD genetics whose phenotype emerges only in the context of a higher-order TF interaction, emphasizing the state dependence of polygenic-to-candidate-core convergence and demonstrating the value of combinatorial perturbations in resolving disease-associated cell states.

## DISCUSSION

A perturbation can reveal gene function through its marginal effect, but it can also create regulatory conditions that reveal the functions of other genes and elements. Our results show that these two forms of utility need not coincide. The magnitude of a TF’s global chromatin response was only weakly correlated with the number of AMD credible sets revealed (*r* = 0.12), and CREB3L3 had little effect on disease-associated programs until combined with FOSB and SOX18. These findings motivate selecting perturbations for the regulatory conditions they create, as well as for their individual effects.

The importance of regulatory state is well established in human genetics.^16, 68, 69^ Here, elements made accessible by TF activation contained 28% of fine-mapped AMD variants, compared with 5 to 7% in resting RPE-1, even though the TFs were selected without knowledge of AMD genetics. Synthetic regulatory perturbations can thus expose substantial disease-associated signal in vitro without prior knowledge of the mechanisms or physiological conditions that surface it in vivo. These perturbation-induced chromatin maps may provide functional priors for fine-mapping^70–72^ and strengthen approaches for enhancer-gene linking.^11, 54, 55^ These benefits extended from regulatory elements to downstream genes: despite using features from a single cellular system, Perturb-PoPS recovered MRD genes from common-variant association signals more effectively than either conventional PoPS or a model trained on unperturbed RPE-1 cells.^57^ Applying these approaches in other disease models could help connect unresolved associations to molecular mechanisms.

Systematically varying regulatory state also revealed how disease-relevant genes respond to combinatorial regulation. Genes nominated directly by AMD GWAS showed no elevated sensitivity to non-additive TF regulation, whereas MRD genes and functionally informed GWAS-prioritized genes identified by PoPS were unusually sensitive. Their shared enrichment in perturbation-derived programs revealed convergence between genes implicated by Mendelian genetics and those prioritized from common-variant associations. These patterns are consistent with the omnigenic model, in which common variation affecting peripheral genes influences disease through regulatory effects on core genes.^30, 31^ Mendelian disease genes have been proposed as empirical proxies for core genes^32, 33^, and their sensitivity to TF combinations and recovery by Perturb-PoPS support this interpretation in our system. Importantly, different TF combinations engaged distinct subsets of these candidate core genes, suggesting that the regulatory relationships underlying this convergence vary with cell state. The broad convergence proposed by the omnigenic model may therefore reflect a superposition of more selective, state-dependent routes to core genes.

This state dependence suggests designing experiments around both disease-relevant genes and regulatory inputs that can reveal their latent function. Chromatin priming provides one criterion for selecting combinations from single-perturbation measurements: pairs in which one TF opened binding sites for the other were more likely to interact neomorphically. Mosaic extended this logic quantitatively, and local chromatin features around each gene were most informative for predicting responses to the most neomorphic TF pairs (*r* = 0.466). Together, these results support using single-perturbation measurements to prioritize informative combinations capable of inducing new regulatory states. Model interpretation extended this search beyond the perturbations used for training: attribution for the AMD convergence program nominated CREB3L3, a regulator outside the pairwise screen. Adding CREB3L3 to FOSB and SOX18 perturbations enhanced the program beyond the expectation from their lower-order effects and induced additional MRD genes absent from the training data. Model-guided experiments can therefore expand the gene responses available for subsequent modeling. These findings suggest an iterative genetics-guided Map-Perturb-Predict strategy for higher-order discovery^73^: map regulatory changes with Multiome Perturb-seq, perturb TFs in combinations nominated by those maps to identify programs supported by human genetics, and predict additional regulators by modeling the responses of those programs.

Emerging genome-wide Perturb-seq datasets^5, 12^ could make this strategy particularly useful in systems where human genetics has already implicated enough genes to define disease-relevant programs. Single-perturbation expression maps could identify panels of regulators that alter diverse subsets of Mendelian and functionally prioritized GWAS genes, providing complementary backgrounds in which to test disease-relevant genes and variants without requiring initial combinatorial screens. Where single-perturbation Multiome maps are available, disease genetics could also guide the choice of regulatory backgrounds and subsequent TF combinations. Perturbations could be prioritized for exposing elements harboring risk variants, while the priming criterion could be refined to favor opening sites for partner TFs at those elements or at elements linked to genetically prioritized genes. This would focus the search on combinations whose disease relevance is not yet apparent from individual transcriptional responses. Extending the approach to systems with less genetic evidence, or beyond TF perturbations, will require further principles for identifying informative regulatory states. Learning to construct such states in vitro would bring more gene functions and disease mechanisms within experimental reach.

## LIMITATIONS OF THE STUDY

The extent to which perturbations can reveal all quiescent variants or relevant gene programs for a trait is limited by sequencing depth, number of perturbations, and perturbation modality. Future work will be required to identify the perturbations that are the most disease informative as our study was limited to relatively unbiased activation of TFs. Studies in other trait-cell type systems will be essential to establish the generalizability of this approach. Our current version of Perturb-PoPS applies minimal feature engineering to capture gene programs from Perturb-seq; future work will be required to maximally extract disease-informative gene programs from Perturb-seq data. Our omnigenic model interpretation cannot draw direct links from polygenic disease genes to monogenic candidate core genes; future work geared at perturbing disease genes alongside TFs will be required to further interpret our findings. Nomination of out-of-distribution TF combinations by Mosaic is limited by chromatin features that implicate families of regulators with similar cognate motifs. In our TF triples screen, we were able to mitigate this either by focusing on unique motifs that suggest a single regulator (CREB3L3) or using orthogonal information to select a top candidate (EGR3). Discovery of latent regulators is also constrained by the fixed motif dictionary used to featurize peaks; future work could extend the modeling framework to operate directly on accessible sequence, allowing de novo motif discovery.

## ACKNOWLEDGEMENTS

We gratefully acknowledge all members of the Norman and Dey labs for valuable thoughts and discussion related to this work. We acknowledge the use of the Integrated Genomics Operation Core (RRID: SCR_027801) and Flow Cytometry Core Facility (RRID: SCR_021105), funded in part by the NCI Cancer Center Support Grant (CCSG, P30 CA008748). Research reported in this publication was supported by NIH grants DP2GM140925 (T.M.N.), R01HG014186 (T.M.N.), U01HG012103 (T.M.N), R00HG012203 (K.K.D.), R01HG014008 (K.K.D.), and F31HG014410 (E.M.). The content is solely the responsibility of the authors and does not necessarily represent the official views of the National Institutes of Health. K.K.D. is supported by the Josie Robertson Investigator Program. K.M.S. is supported by the Alan and Sandra Gerry Metastasis and Tumor Ecosystems - GMTEC Metastasis Scholars Fellowship. R.C.A. is a Robert Black Fellow of the Damon Runyon Cancer Research Foundation (DRG-2462-22). Part of this research has been conducted using the UK Biobank Resource under Application Number 16549.

## AUTHOR CONTRIBUTIONS

Conceptualization, E.M., K.G., K.M.S., K.K.D., and T.M.N.; Methodology, E.M., K.G., K.M.S., K.K.D., and T.M.N.; Software, E.M. and K.G.; Formal Analysis, E.M., K.G., K.K.D., and T.M.N.; Investigation, E.M., K.M.S., and R.C.A.; Visualization, E.M. and K.G.; Writing – Original Draft, E.M., K.G., K.K.D., and T.M.N.; Writing – Review & Editing, E.M., K.G., K.K.D., and T.M.N.; Supervision, K.K.D. and T.M.N.; Project Administration, K.K.D. and T.M.N.; Funding Acquisition, K.K.D. and T.M.N.

## RESOURCE AVAILABILITY

Processed single-cell data for Multiome and combinatorial Perturb-seq screens will be deposited to Zenodo upon publication. All custom analysis code generated in this study will be deposited to Zenodo and made available on Github upon publication. Plasmids generated in this study will be made available on Addgene upon publication. Requests for further information and resources should be directed to and will be fulfilled by the lead contact, Thomas M. Norman.

## SUPPLEMENTAL INFORMATION

**Document S1:** Figures S1-S10, Supplemental Figure legends, and Supplemental Note.

**Table S1:** sgRNA protospacers used in this study.

**Table S2:** Differential motifs for each perturbed subpopulation in the Multiome Perturb-seq screen, calculated via ChromVAR and peak-free scoring.

**Table S3:** Global interaction metrics and motif cross-regulation for 1,225 TF pairs profiled in the combinatorial Perturb-seq screen.

**Table S4:** Mendelian, PoPS, and MAGMA retinal disease genes.

**Table S5:** Gene features, weights, and perturbation loadings for 100 transcriptional programs generated via non-negative sparse PCA applied to the combinatorial Perturb-seq screen.

**Table S6:** Oligonucleotides used in this study.

## DECLARATION OF GENERATIVE AI AND AI-ASSISTED TECHNOLOGIES IN THE MANUSCRIPT PREPARATION PROCESS

During the preparation of this work, the authors used Claude (Anthropic) and ChatGPT (OpenAI) to improve readability, style, and conciseness. After using these tools, the authors reviewed and edited the content as needed and take full responsibility for the content of the published article.

## METHODS

### Experimental

#### Cell lines and lentiviral packaging

RPE-1 cells stably expressing the dCas9-VPH CRISPRa effector, generated in our previous work^40^, were maintained in DMEM:F12 (Gibco, 11320033), 10% Fetal Bovine Serum (FBS) (VWR, 97068-085), 100U penicillin and 100 µg/mL streptomycin (Gibco, 15140122), and 0.01 mg/mL Hygromycin B (Thermo Fisher, 10687-010). LentiX 293T cells were purchased from Takara Bio (#632180) and maintained in DMEM (Gibco #11965092), 10% Fetal Bovine Serum (FBS) (VWR #97068-085), 100U penicillin and 100 mg/mL streptomycin (Gibco, 15140122). For lentiviral packaging, LentiX cells were grown to 80% confluence and co-transfected with library vector and packaging plasmids psPAX2 and pMD2G using TransIT-LTI Transfection Reagent (Mirus #MIR2300). 24 hr post-transfection, media was changed to DMEM + 10% BSA (Sigma #A9418). Virus was harvested by filtering collected media through a 0.45µm filter 48 hr post-transfection. Titers of sgRNA-containing lentivirus were quantified by transducing 3 × 10^5^ RPE-1 VPH cells with varying amounts of lentivirus in a 6-well plate and measuring GFP positivity by FACS 3 days post-transduction.

### Multiome Perturb-seq: sgRNA library construction and quality control

Vector pEM070 (Fig. 1A, top) was created from pEM040 (Addgene #224898) by digesting with BsiWI and BbsI, removing the eGFP coding sequence, and replacing with a gBlock (IDT) encoding eGFP with a C-terminal fusion of the KASH domain from human SYNE2 (isoform 1, amino acids 6804-6885) via HiFi assembly (NEB #E2621). To construct the Multiome single-TF library, pEM070 was digested with BstXI and BlpI to remove the LacZ stuffer, and the 8377 bp backbone was purified from a 0.8% agarose gel. 141 sgRNAs against 120 TFs were chosen from our previous TFome-wide CRISPRa screen^40^, using strength of on-target activation, lack of “bad seed” protospacer motifs, number of upregulated genes with promoters inaccessible in resting RPE-1 cells (a transcriptome-only proxy for pioneering activity), and expression in the human retina as selection criteria. 21 TF targets with uniquely strong transcriptional phenotypes were targeted with two sgRNAs, and the remaining 99 were targeted with one sgRNA. 16 unique non-targeting control sgRNAs were selected, for a total library size of 157 sgRNAs. The library was ordered as an Oligo Pool (Twist) with appropriate 5’ and 3’ overhangs for assembly into pEM070. The pool was resuspended per manufacturer instructions at 5 ng/µL, and 10 ng was amplified via 5 cycles of PCR with primers P276F and P276R (Table S6) using 2x KAPA HiFi Master Mix (Roche #07958935001). The double-stranded PCR product was purified (NEB #T1030) and cloned into the digest of pEM070 via HiFi Assembly (NEB) for 1 hr at 50°C, using a 5:1 molar ratio of library insert to vector. Transformation was performed with an Eppendorf Eporator into Endura Electrocompetent Cells (LGC60242-2) per manufacturer protocol. Transformants were incubated in 1 mL Recovery Media at 32°C for 1.5 hr, then expanded into 100 mL LB + carbenicillin for overnight culture. Plasmid libraries were produced via midiprep (Qiagen #12941).

For QC of the constructed library, sgRNA protospacers were amplified via 2x KAPA HiFi Master Mix from the above midiprep with primers containing Illumina P5 and P7 adapters and Nextera i7 indices (oJR232/oJR233, Table S6). Following 2x Pronex purification (Promega #NG2001), amplicons were sequenced on an Illumina MiSeq with PE70 read structure. The resulting fastq files were processed with a custom Python script to quantify representation of each sgRNA in the library.

### Dual-guide Perturb-seq: sgRNA library construction and quality control

Vector pRCA672 (pDCG-SFFV-GFP) was derived from pHAGE-EF1aL-eGFP-W (Addgene #126686) via HiFi assembly (NEB #E2621) of the following fragments: guide cassettes, amplified from pBA904 (Addgene #122238); SFFV promoter, amplified from pHR-UCOE-SFFV-Zim3-dCas9-P2A-BFP (Addgene #188767); and GFP coding sequence, amplified from pRCA360 (Addgene #238164). For library cloning, pRCA672 was digested with BstXI and BlpI, and the backbone was purified from a 0.8% agarose gel. 50 of the 141 targeting sgRNAs screened via Multiome Perturb-seq were chosen for pairwise transcriptome profiling based on strength of ATAC and motif enrichment phenotypes. Targets with representation of less than 20 cells in the Multiome screen were not considered due to sparsity of ATAC profiles. To ensure even representation in the dual-guide screen, we estimated fitness effects by comparing each sgRNA’s initial abundance in the Multiome library to its cell number in the Multiome screen. We then ordered targeting libraries for “A-position” and “B-position” sgRNAs as Oligo Pools (Twist), accounting for fitness effects by ordering one copy of each sgRNA with fitness advantage (e.g., MYC); two copies of sgRNAs with minimal changes in fitness (e.g., IRF4); and four copies of sgRNAs with apparent fitness defects (e.g., GATA2). PRDM16 was removed due to its abnormally large positive fitness effect, which we reasoned would result in a highly unbalanced final cell population. Differential genes and peaks compared to all control cells in the Multiome screen were called for each NTC sgRNA, and the sgRNA with the fewest differential features was selected as the control sgRNA for single perturbations (targeting sgRNA + control sgRNA) with four copies in each library. Separate A-position and B-position Oligo Pools containing one copy each of 9 NTC sgRNAs, similarly selected from the Multiome screen, were also ordered to construct a control-only library.

Targeting and control library amplicons were generated via PCR stitching of A-position and B-position Oligo Pools as follows. First, 10 ng of each Oligo Pool was amplified via 8 cycles of PCR with NEBNext Ultra II Q5 (NEB #M0544) using primers oDOS384 and oDOS531 (A position) or oDOS534 and oDOS535 (B position). An hU6 promoter fragment was similarly amplified from vector pJR98 (Addgene #187239) with oDOS532 and oDOS533, adding overhangs with the A-position and B-position amplicons. After purification (NEB #T1030), equimolar amounts of the three amplicons (2.129 pmol, 100 ng of the smallest fragment) were stitched together via PCR with KAPA HiFi Master Mix, per the following program: 95° 3’, 5 cycles of [98° 10”, 65° 15”, 72° 10”], 72° 2’, 4° hold. The reactions were placed directly on ice, and 15 pmol of primers oDOS384 and oDOS535 were added to selectively amplify full-length stitched product with the following program: 4° 5’, 95° 3’, 5 cycles of [98° 10”, 65° 15”, 72° 10”], 72° 2’, 4° hold. Full-length stitching products, containing all pairwise combinations of A-position and B-position library elements, were run on an E-Gel EX 1% (Thermo #G401001) to verify size and purified via 1.2x Pronex. Purified products were each cloned into digested pRCA672 via HiFi Assembly. Targeting and control plasmid libraries were then produced via electroporation, overnight culture, and midiprep as above. For quality control, sgRNA protospacer cassettes (containing the A-position sgRNA on the 5’ end and the B-position sgRNA on the 3’ end) were amplified from plasmid libraries via 12 cycles of PCR with NEBNext Ultra II Q5, purified via 1.15x Pronex, and sequenced in a PE50 run on a NovaSeq 6000 (Illumina) using the S1 reagent kit at a loading concentration of 0.8-1.2 nM with 1% PhiX spike-in. During virus production, the control plasmid library was spiked into the targeting library at a molar concentration of 1%, and LentiX cells were transfected with the combined library pool.

### Multiome Perturb-seq screen

On Day 0 of the screen, 2.4 × 10^7^ RPE-1-VPH cells were transduced with combined library lentivirus at 250 µL unconcentrated lentiviral supernatant / million cells and 8 µg polybrene / mL media. This population was expanded in 6 T175 flasks and sorted for GFP positivity on Day 3. 3.2 × 10^5^ cells were recovered from sorting (titer of 0.9%) and plated in one T25 flask. The sorted population was expanded for 13 additional days. On Day 16, 2 × 10^6^ cells were processed to isolate nuclei, following the 10x protocol “Nuclei Isolation for Single Cell Multiome ATAC + Gene Expression Sequencing” (#CG000365, Rev C) with 4 min lysis time (Step 2d). Nuclei were resuspended in 500 µL PBS + 1 U/µL Sigma Protector RNase inhibitor (Sigma #3335402001), and 25 µL 7-AAD (Thermo #00-6993-50) was added to stain intact nuclei. Samples were immediately sorted by FACS (BD FACSAria) to purify 7-AAD / GFP double-positive nuclei and remove debris. Sorted nuclei were resuspended in Nuclei Buffer targeting a concentration of 4,000 nuclei/µL. 18,000 nuclei were loaded into each transposition reaction per manufacturer protocol (Chromium Next GEM Single Cell Multiome ATAC + Gene Expression User Guide, #CG000338 Rev F) and encapsulated across 6 GEM groups with the Chromium Controller, Chromium Next Gen Chip J, and Chromium Next GEM Single Cell Multiome ATAC + Gene Expression Reagent Bundle (10x Genomics #1000283) with no protocol modifications. We used 7 index PCR cycles for ATAC library amplification (Step 5.1d); 6 PCR cycles for cDNA amplification (Step 6.1d); and 12 PCR cycles for Gene Expression library amplification (Step 7.5d). ID libraries were generated via T7-assisted amplification from cDNA as previously reported^18^, with 7 index PCR cycles. After PicoGreen quantification and quality control by TapeStation (Agilent), libraries for each GEM group were pooled by molarity and sequenced on a NovaSeq X (Illumina) in a PE28/88 run with a target sequencing depth of 20,000 reads/cell (GEX libraries) or 4,000 reads/cell (ID libraries); or in a PE150 run with a target sequencing depth of 25,000 reads/cell (ATAC libraries). The loading concentration was 0.8-1.2 nM and a 1% PhiX spike-in was added for quality control.

### Dual-guide Perturb-seq screen

On Day 0, 2.4 × 10^7^ RPE-1-VPH cells were transduced with 180 µL combined library lentivirus / million cells and expanded as above. 1.2 × 10^6^ GFP+ cells were recovered after sorting on Day 3 (titer of 5.5%) and plated in one T75 flask for 13 additional days of outgrowth. On Day 16, cells were washed and resuspended in PBS + 1% BSA (Sigma #126615). The concentration of GFP+ single cells was measured via FACS (Thermo Attune) and adjusted to 2,000 cells / µL with addition of PBS/BSA. This cell suspension was loaded into each RT reaction, targeting 2.5x overloading compared to the maximum supported 20,000-cell recovery target, and cells were encapsulated across 8 GEM groups on the Chromium X using the Chromium GEM-X Single Cell 3’ Chip Kit (10x Genomics #1000690) and GEM-X Universal 3’ Gene Expression v4 Reagents (10x Genomics #1000691, Protocol #CG000731 Rev B) with no modifications.

Although sgRNAs are captured and barcoded via Capture Sequence 1 on GEM-X 3’ beads, CRISPR libraries are not officially supported in the GEM-X 3’ v4 chemistry, so we modified the cDNA amplification protocol as follows. The GEM-X cDNA Primers (#2000089) were replaced with Feature cDNA Primers 1 (#2000096) from the Next GEM 3’ Feature Barcode Kit in Step 2.2a, and cDNA was amplified with 10 PCR cycles (less than the recommended 11 given overloading). In the cDNA cleanup (Step 2.3), the supernatant in Step 2.3d containing the barcoded sgRNAs was retained. After the cDNA for Gene Expression libraries was generated (Step 2.3n), an additional 1.2x SPRIselect was performed on the Step 2.3d supernatant. Gene Expression libraries were generated from the Step 2.3n product with 8 index PCR cycles. CRISPR libraries were generated from the purified Step 2.3d supernatant following Step 4 of the Chromium Next GEM Single Cell 3’ v3.1 User Guide (#CG000316 Rev E). Gene expression and CRISPR libraries for each GEM group were quantified via PicoGreen and visualized via TapeStation for quality control, then pooled by molarity and sequenced on a NovaSeq X in a PE28/88 run with a target sequencing depth of 20,000 reads/cell (GEX libraries) or 5,000 reads/cell (ID libraries). The loading concentration for each library was 0.8-1.2 nM, and a 1% PhiX spike-in was added for quality control.

### TF triples screen

Vector pEM125, an sgRNA expression vector with mCherry fluorescent marker, was created by digesting pRCA672 with NcoI and SbfI (NEB) and replacing the GFP marker with a PCR amplicon of the mCherry coding sequence via HiFi Assembly. Single sgRNAs were ordered from IDT as forward and reverse oligos, then annealed (Table S6): sgFOSB, sgSOX18, sgIRF4, and sgNTC were selected from the dual-guide Perturb-seq experiment; sgCREB3L3 and sgEGR3 were chosen from Southard*, Ardy* et al.^40^ on the basis of highest on-target activation (CREB3L3) or strongest effect on Program 38 (EGR3). pEM125 was digested with BstXI and BlpI, and annealed sgRNA oligos were cloned into the digest via T4 ligation (NEB #M0202). The FOSB_SOX18 sgRNA cassette was ordered as a Gene Fragment (Twist) and cloned into a BstXI/BlpI digest of pRCA672 via HiFi assembly. All constructs were transformed into NEB Stable Competent E. coli (#C3040H) and miniprepped for transfection (NEB #T1110). Lentivirus encoding each individual construct was produced as above. On Day 0 of the screen, 3.0 × 10^5^ RPE-1-VPH cells were transduced in 6-well plates with unconcentrated lentivirus encoding individual single-sgRNA constructs, with or without the FOSB_SOX18 dual-sgRNA construct, using 8 µg polybrene / mL media. On Day 3, each condition was sorted to select for mCherry/GFP double-positives (FOSB_SOX18 conditions, including a double-only condition where the single expression vector encoded a control guide) or mCherry positives (single-sgRNA and NTC conditions). Cell populations were expanded until the end of the screen and passaged when 80% confluent. On Day 16, cells were washed with cold PBS and lysed in wells via addition of 500 µL DNA/RNA Shield (Zymo #R1100). Lysate was sent in 3 technical replicates per condition to Plasmidsaurus for bead-based total RNA extraction, oligo-dT-primed reverse transcription, second strand synthesis, and cDNA library construction. Sequencing was performed on an Illumina NovaSeq X Plus targeting 20 million single-end reads per replicate.

### Computational

#### Use of AI tools in computational analysis

Software for analysis and visualization generated in this study was developed, maintained, documented, and debugged in part by Claude (Anthropic), ChatGPT (OpenAI), and Gemini (Google). All AI-generated code was edited and tested before use in analysis workflows.

### Multiome Perturb-seq: data processing and quality control

Gene Expression and ATAC fastq files were aligned to the transcriptome and genome respectively (reference: cellranger-arc-GRCh38-2020-A-2.0.0) with cellranger-arc-2.0.2 (10x Genomics), using cellranger-arc count with default parameters. IDs were assigned to cells as previously reported^18^, using a Top ID log UMI threshold of 3 and HDBSCAN epsilon of 0.5. QC metrics were calculated with scanpy, and the population was filtered to cells assigned exactly one sgRNA and passing UMI, GEX unique genes, ATAC unique fragments, and ATAC TSS enrichment thresholds (Fig. S1). The GEX matrix was processed by removing mitochondrial genes and genes expressed in less than 6% of cells, then z-normalizing counts to GEM group controls using the perturbseq codebase^17^ and regressing out cell cycle phase. The ATAC matrix was filtered to 250,000 tile features using SnapATAC2, and peaks were called with MACS3, grouping both by sgRNA and by TF target. Peak matrices were quantified with paired-insertion counting. For downstream analyses, populations (sgRNAs or target TFs) containing fewer than 20 cells were removed.

### Multiome Perturb-seq: descriptive analyses

Spectral embedding (SnapATAC2^79^) was performed on the Multiome ATAC tile matrix resulting in 30 spectral components. The first spectral component was highly correlated with ATAC library size and removed for downstream analysis. PCA was performed on the Multiome GEX matrix with 29 components to match the dimensionality of the spectral embedding. To generate the joint UMAP in Fig. 1D, embeddings were pseudobulked by averaging across sgRNAs. Components in both pseudobulk embeddings were scaled to zero mean and unit variance, and rows were L2-normalized, such that neither embedding “outweighs” the other based on differences in scale alone. Canonical correlation analysis (scikit-learn, 29 components) was then performed on the normalized embeddings, and the resulting transformed matrices were concatenated. UMAP (metric = ‘correlation’, n_neighbors = 10) was then fit to the concatenated CCA components. In the resulting scatterplot, each perturbed subpopulation is colored by target TF on a continuous scale in alphabetical order such that homologs (e.g. FOXA1, FOXA3, FOXC1, FOXC2, …) are colored similarly.

To determine the strength of transcriptome vs. ATAC phenotypes, energy distance was calculated in embedding space for each modality, again using 29 GEX PCA components and 29 ATAC spectral components. Each single-cell embedding matrix was again L2-normalized to place distances on a common scale; this projects each cell’s embedding vector onto the unit sphere, such that the subsequent distance calculations compare each cell’s “direction” in embedding space relative to control cells. Energy distance was calculated using the dcor library with exponent = 2 (squared Euclidean metric) and estimation_stat = ‘U’ (unbiased) as previously reported.^43^ To further avoid confounding by cell number, a fixed sample of 200 control cells, around the average representation of cells per target TF, was used for each calculation. Significance was assessed over 1,000 permutations of cell labels.

Differential ATAC peaks were called via Mann-Whitney tests with Benjamini-Hochberg correction on the single-cell peak matrix (grouping cells by target TF) and labeled with their nearest gene. To identify high-confidence newly expressed genes (Fig. 1F), we first calculated percentile expression changes between control and perturbed cells for each target TF across all genes not present in the filtered GEX matrix, as previously reported.^40^ Differential peaks were matched to newly expressed genes, and genes detected in at least 10% of cells with a percentile expression change of at least 10 and at least one concordant (opening) differential peak were retained as high-confidence newly expressed genes.

### Motif enrichment and determination of “primed” TF pairs

Motif enrichment was performed via the GPU-accelerated implementation of ChromVAR available in the scprinter library^80^ and via a custom peak-free single-cell scoring method (scripts available on GitHub upon publication). For ChromVAR, peaks were called by sgRNA, and the background was sampled from control cell peaks. Deviations for 1,019 JASPAR2026 motifs^81^ were computed in each cell and z-normalized to control cells. Differential motifs in each subpopulation compared to control cells (grouping by target TF) were called via a Kolmogorov-Smirnov test on z-normalized deviations, and FDR-corrected *q* values were generated by the Benjamini-Hochberg procedure. For peak-free single-cell motif scoring, genome-wide coordinates for 827 available JASPAR motifs were identified by downloading motif tracks from the UCSC Genome Browser and filtering to matches with score ≥ 300 to generate .bed files with coordinates for each motif occurrence (`download_motifs.py`, `merge_motifs.py`, `bed_motifs.py`). Single-cell .bam files were created from each GEM group’s ATAC library with samtools split on the ‘CB’ tag (`sc_bams.py`), and bigwig files were created from each single-cell .bam with deepTools bamCoverage (`sc_bws.py`). Coverage for each single cell across each of the genome-wide motif tracks was then calculated with bigWigAverageOverBed using each motif’s .bed file (`sc_motif_scores.py`). Raw coverage scores were z-normalized to control cells, and differential motifs were identified as above.

To identify “primed” pairs of TFs, we adopted a simple definition based on motif co-enrichment. We first assigned on-target motifs to 48 of 50 TFs in the dual-perturbation screen (MYRF and ARID1A have no motifs in JASPAR2026), using PWMs from mouse if no human data were available. We took the union of differential motifs from ChromVAR and peak-free scoring, keeping the larger z-score as the effect size if a motif was detected by both methods. We categorized a TF pair AB as “primed” if either activation of A caused opening (differential with positive z-score) of B motifs, or activation of B caused opening of A motifs. A primed pair was further considered “bidirectional” if activation of A opened B motifs and activation of B opened A motifs. Priming of pairs containing MYRF or ARID1A was determined exclusively by enrichment of partner motifs upon activation of MYRF or ARID1A, and none of these pairs were considered bidirectional. From this list, we excluded primed pairs belonging to the same TF family with substantially similar motifs based on JASPAR clustering, reasoning that in these cases we likely cannot differentiate between, e.g., FOXA3 opening a FOXA1 motif and FOXA3 opening its own motif. Our grouping of TF families is shown in Fig. 1H (smaller spacing between sectors of the circos plot). Some related TFs (e.g., NKX2-5 and NKX3-1) have motifs which are clearly dissimilar based on visual inspection of PWMs, so these pairs were maintained as primed if their motifs were co-enriched. We also excluded rare instances where activation of A alone was sufficient to induce expression of B (or vice-versa), reasoning that co-enrichment in these cases was likely due to a *trans* effect rather than true cooperation within the epigenome. To construct the circos plot in Fig. 1H, a single chord was drawn between the TFs of a primed pair and colored by the TF whose motif is opened. Chords between bidirectional pairs are colored by the TF whose motif has the larger z-score.

### Dual-guide Perturb-seq: Data processing and quality control

Gene expression fastq files were aligned to the transcriptome with cellranger-9.0.1 (10x Genomics), using cellranger count with default parameters. Expression matrices were aggregated and initial ID calls made with cellranger aggr (--normalize none). To supplement and assess the consistency of cellranger’s ID calls, we set thresholds at the maximum of 3 UMI and the 25^th^ percentile of each ID’s UMI distribution across all cells, and we assigned an ID to a cell if that cell contained more ID UMIs than its corresponding threshold. We made assignments by each procedure independently and retained cells assigned exactly one A-position and one B-position feature in either case. In rare instances where cellranger’s assignment was not concordant with manual assignment, we kept the cellranger assignment; if cellranger made no assignment, we kept the manual assignment. Orientation-aware assignments (e.g., AB and BA each maintained as separate subpopulations, 50 × 50 = 2,500 possible pairs) were maintained for quality control analyses in Fig. 2A and Fig. S2C. For downstream analyses, orientations were merged and each assignment was labeled unambiguously with single perturbations in alphabetical order, leaving 1,225 (50 choose 2) possible pairs. QC metrics were calculated, and the assigned cell population was filtered based on thresholds for total UMI and unique genes (Fig. S2B). Genes expressed in less than 5% of cells were removed, and the remaining genes were z-normalized to GEM group controls. PCA (50 components) was fit to the expression matrix, and energy distance to control cells for all 50 single perturbations and 1,210 of 1,225 double perturbations with at least 20 cells (Fig. 2B) was calculated on L2-normalized principal components as above. Energy distance distributions between single and double perturbations were compared with a one-sided Mann-Whitney test. Differentially expressed genes were called using a Kolmogorov-Smirnov test on z-normalized expression values and Benjamini-Hochberg correction (perturbseq), grouping by target with orientations merged (1,275 total groups, 1,225 pairs + 50 singles).

### RPE / choroid single-cell atlas analysis

An AnnData file containing snRNA-seq profiles of 566,546 human RPE, choroid, stromal, and vascular cells was downloaded from the Cell Annotation Platform of the Human Cell Atlas (v3, June 10^th^, 2026).^44^ Counts were normalized and log-transformed with scanpy following standard preprocessing pipelines. Highly variable genes were selected and marker genes for each cell type were identified (grouping by “author_cell_type”) with scanpy’s `rank_genes_groups()` function (method = ‘wilcoxon’), comparing each cell type to the rest of the dataset and filtering to genes specifically upregulated in each state with q *<* 0.1. The z-normalized expression matrix from the dual-guide experiment was pseudobulked by perturbation target, and for each of the five RPE subtypes labeled as distinct cell types, state scores for each target (singles and doubles) were calculated as the dot product of the target’s mean expression of marker genes present in the normalized matrix and these marker genes’ specificity scores generated by `rank_genes_groups()`. Scores within each state were z-normalized across all singles and doubles for plotting, and variance of singles and doubles was compared via Levene’s test. The five resulting p-values were corrected for multiple hypothesis testing via Benjamini-Hochberg.

### Characterization and quantification of genetic interactions

Linear models regressing double transcriptional profiles on profiles of their respective single perturbations were fit as previously reported.^17^ Briefly, perturbation target genes and genes with mean UMI less than 0.25 were removed from the z-normalized expression matrix. Robust linear models (statsmodels) using Tukey’s Biweight M-estimator were fit across genes to predict each double perturbation’s z-normalized expression as a function of the z-normalized expression of each of its single perturbations.

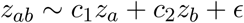

To characterize interactions while avoiding measurement noise at the level of single genes, we first fit a 64-component scVI embedding^46^ to the raw count matrix, taking these coordinates to represent each cell’s transcriptional state. We pseudobulked by target and normalized by subtracting the control cells’ embedding from that of all other targets, then performed a whitening procedure to prevent large axes of variation (e.g., cell cycle, EMT) from overwhelming the transcriptional heterogeneity in the dataset while compressing axes of variation that were likely due to single-cell sampling noise. We performed singular value decomposition on the NTC-normalized embeddings, resulting in 64 orthogonal components. We then partitioned all single cells into four disjoint splits, stratifying within each perturbation such that every split contained a representative sample of cells. Within each split, we again pseudobulked by target to estimate the original scVI embedding and projected these estimates onto the SVD components. The variance of each component across each of the four splits gives an estimate of sampling noise, and we took the median variance across all 64 components as the “noise floor” in the experiment after scaling to account for the size of each split relative to the dataset. We then whitened the embedding matrix by projecting the original pseudobulk embeddings onto the SVD axes and divided each component’s coordinates by either the square root of its variance or the noise floor, whichever was larger. Using this whitened perturbation-effect matrix, we calculated statistics that quantified interaction nonlinearity (*neomorphic score*) and synergy (*synergy score*), and *similarity* to single perturbations as follows.

### Neomorphic score

We define a *neomorphic interaction* as a double phenotype AB that cannot be reconstructed as a linear combination of single phenotypes A and B. To quantify the strength of neomorphic interaction across pairs, we resolved each embedding AB into an additive component within the plane spanned by singles A and B and a neomorphic component orthogonal to this plane. The embedding of a pair AB whose interaction phenotype is a linear combination of singles A and B will lie within or close to the plane spanned by A and B, producing a small neomorphic component. In contrast, if A and B interact nonlinearly, AB will be embedded outside this plane and produce a large neomorphic component. In this model, then, the magnitude of AB’s neomorphic component corresponds to the strength of the nonlinear interaction between A and B. We calculated an orthonormal basis for the plane spanned by A and B by stacking the embeddings of A and B into a 64 × 2 matrix and performing QR decomposition. We then projected the embedding of AB onto this basis to derive the additive component and subtracted this from the original embedding of AB to find the neomorphic component.

Raw neomorphic components were negatively correlated with the cell number of each double. To “regress out” this effect, we compared each double’s neomorphic component to that of other doubles with similar cell-number characteristics. For each double, we calculated 1) its log1p-transformed cell number, 2) the log1p-transformed harmonic mean of the cell number of its two singles, and 3) the absolute value of the log-transformed ratio between the cell numbers of each of its singles. We z-normalized each feature across all doubles, then identified the 100 nearest neighbors for each double in this standardized feature space not sharing either single, such that correction for a given double only considers pairs with no common perturbations. After weighting neighbors by a Gaussian kernel on their distance in cell-count feature space, we computed a weighted median and median absolute deviation of the log-transformed raw neomorphic component across that double’s 100 neighbors. Each double’s neomorphic score was then calculated as the difference between its log-transformed raw neomorphic component and the weighted median of its cell-count-matched neighbors, scaled by the robust standard deviation of its neighbors (local MAD scaled by a normalizing constant to approximate a standard deviation).

### Synergy score

Consistent with previous literature, we define an interaction as *synergistic* if the magnitude of the double phenotype is greater than the additive expectation of its single phenotypes and *buffering* if its magnitude is less than the additive expectation. Since we set the origin of the latent space to the embedding of control cells, lengths of embedding vectors correspond to phenotype magnitudes. To quantify synergy, we take the difference between the additive component of each pair AB and the additive expectation of singles A and B. We only consider the additive component here to avoid double-counting the neomorphic component of each interaction, which by construction is orthogonal to single phenotypes A and B. To fit the additive expectation, we regressed the L2 norm of double embeddings on that of the vector sum of single embeddings. The synergy score for each double was then calculated as the difference between the norm of its additive component and the scaled norm of the vector sum of its singles (multiplying by the coefficient derived from the regression).

### Similarity

Similarity of each double’s transcriptional profile to that of each of its singles was simply calculated as the cosine similarity in embedding space. The single with greater cosine similarity to each double was labeled as that double’s “dominant target.”

### Generation of gene programs by sparse PCA

We fit sparse PCA programs to the gene expression matrix as previously reported^40^ with slight modifications. Briefly, the z-normalized and filtered single-cell expression matrix was pseudobulked by target. Genes were subset to those which were 1) differentially expressed in at least one perturbed subpopulation with *q <* 0.01 and |*z*| *>* 0.25; and 2) members of the PoPS gene universe.^57^ We next masked each target gene’s expression by setting its z-score to 0 within the subpopulations where it was targeted (for example, in the row corresponding to the JDP2_FOSB population, features JDP2 and FOSB were set to 0; in any other subpopulation not targeting JDP2 or FOSB, their z-normalized expression values were left unchanged). This approach allows for cross-regulation among our TF targets to inform the structure of the gene programs, while mitigating large changes in covariance driven exclusively by on-target activation. We then determined the optimal number of programs via the bootstrapping and clustering procedure reported in our previous work.

### Calculation of gene-level residuals

To calculate gene-level interaction residuals that are robust to the noise inherent in scRNA-seq, we adapted a previously reported approach^82^ expressing interaction phenotypes as t-statistics, taking into account both the magnitude (changes in z-normalized expression beyond those detected in singles) and confidence (sizes of the cell populations from which the z-score estimates were derived) of observed phenotypes. We first accounted for saturation effects at extreme values of the observed z-scores by fitting a global linear regression across all double perturbations and non-target genes in the filtered expression matrix, predicting double expression as a separable quadratic function of single expression:

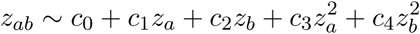

We then calculated raw interaction residuals for each gene and double perturbation by subtracting the expression predicted by this linear model from the observed expression:

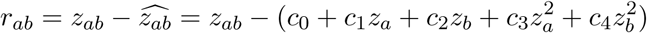

We finally calculated standard errors of the estimates of expression in double AB, single A, and single B, adjusting the variance of singles expression via the delta method to account for the linear model coefficients, and divided the raw residual by their sum to calculate the final residual t-statistic:

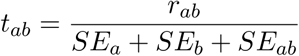

Where 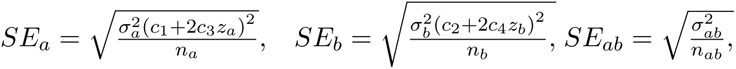, for observed expression variance 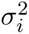 and cell number *n_i_* in subpopulation *i*. For genetics analyses in Fig. 5, the PCI score was calculated for each gene *g* by taking the L_2_ norm of its residual vector ***t_ab_*** across all doubles:

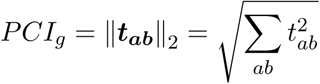

By construction, this score aggregates all deviations from pure additivity, such that higher PCI scores equally reflect regulatory contributions to a gene’s expression from TF pairs exhibiting synergistic, epistatic, and neomorphic responses.

### Modeling residuals from single-TF expression and accessibility

We first constructed a peak-by-feature matrix as follows (`construct_feature_matrix.ipynb`). The Multiome ATAC matrix was filtered to only perturbations present in the dual-guide screen and control cells, and peaks were re-called, grouping by target. Tabular motif features in each peak were generated as previously reported by Fu et al.^60^ Briefly, we used MOODS^83^ to scan the sequence in each peak against 1,019 JASPAR2026 PWMs, returning a binding score for each match. Within JASPAR motif clusters (resolution subcluster_ncor_0.7), sequence matches to multiple PWMs were deduplicated by retaining only the match with the highest score within each cluster, and the resulting peak-by-motif-cluster matrix was MinMax-normalized. We complemented motif features with residualized accessibility values for each of the 50 single perturbations in the dual-guide screen. For each peak, single-cell counts were normalized by each cell’s ATAC library size and aggregated by target. We then subtracted the mean accessibility per peak across all targets from each row, and the mean accessibility per target from all columns, adding back the global mean and scaling each peak’s values to unit variance. The result was a “residualized accessibility” value for each target in each peak. These features (50 targets × 314,512 peaks) were concatenated to the motif features, along with a single “absolute accessibility” feature that simply quantified the normalized read count in each peak across all single cells in the dataset. We then added indicator features for ENCODE CRE identity (.bed file downloaded from ENCODE v3 SCREEN^84^, all human CREs), extending promoter definitions using the Eukaryotic Promoter Database.^85^ TSS annotations for each gene were obtained from ENCODE.^11, 54^ Genes for modeling were filtered starting with those for which residual t-statistics were calculated, then excluding TF target genes, non-protein-coding genes, and genes with no annotated TSS. Each of the remaining 7,780 genes in the modeling universe was mapped to a set of 128 peaks nearest its TSS (pyranges), which was sorted by distance and retained for downstream modeling tasks.

For the bilinear and simple nonlinear models (Fig. S9B), the peak-by-feature matrix was aggregated at gene level to create a gene-by-feature design matrix. Within each gene, the correlation of residualized accessibility with z-normalized expression across all 50 dual-guide targets in the Multiome dataset was calculated for each peak. Within each gene’s 128 peaks nearest to its TSS, features were then aggregated via a correlation-weighted sum, maintaining the sign of the correlation coefficient. This empirically provided better performance than simple averaging or weighting by unsigned (absolute value) correlation with expression. For each gene, we then concatenated 50 single expression features from the dual-guide experiment, resulting in a “cis-chromatin representation” matrix ***X*** of 7,780 genes × 513 features. For the *trans*-chromatin representation, ChromVAR z-scores were aggregated for each single target, and a 32-component PCA embedding was fit to the pseudobulk z-scores. To represent each double perturbation, we combined single PCA components via a permutation-invariant concatenation operation, mapping each double AB to a 64-dimensional vector [*A* + *B, A* × *B*] containing the sum and product of the PCA embeddings of singles A and B. The “*trans*-chromatin representation” matrix ***M*** contains these embeddings for all 1,225 pairs. We then fit bilinear models of the following form:

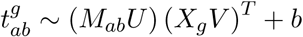

where 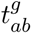 is the residual of gene *g* in pair AB, *M_ab_* is the *trans-*chromatin representation of pair AB, *X_g_* is the cis-chromatin representation of gene *g*, *U* and *V* are learned linear projections with rank 64, and *b* is a learned global scalar bias term. We split genes into 12 cross-validation folds by chromosome as previously reported^86^ and fit the model by gradient descent (AdamW optimizer, learning rate 1e-3, weight decay 1e-3) to minimize the mean squared error between observed and predicted residuals for all training genes. The validation MSE was checked every 1000 steps, and training was stopped if it did not improve after 5000 steps. The model checkpoint with the lowest validation loss was used for inference. The simple nonlinear models were fit similarly, replacing linear transformations *U* and *V* with *f_M_* and *f_X_* parametrized via MLPs with two fully connected layers mapping from the feature spaces of *X* and *M* to hidden dimension 128, then to 64 dimensions:

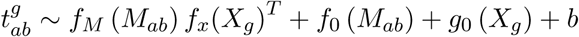

SiLU activation and dropout of 10% were applied between layers of each MLP. Bias terms *f*_0_*, g*_0_ specific to each row of *X* or *M* were learned as linear transformations of the features *M_ab_* or *X_g_*. A global bias *b* was also learned as a single scalar. Training and cross-validation proceeded identically to the bilinear models.

The transformer-style model (Fig. S9A) allows flexibility to consider each peak in the context of all other peaks assigned to a given gene, rather than aggregating peak information via a fixed rule to create one vector for each gene. We therefore learned a function *f* mapping *cis*-chromatin and expression information for gene *g*, as well as trans-chromatin information for singles A and B, to the residual t-statistic for double AB in gene *g*.

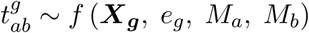

Here, ***X_g_***∈ R^128^*^×^*^464^ is a matrix of each gene’s 128 peaks, sorted by absolute distance to the TSS of *g*, containing motif, accessibility, CRE identity and expression correlation features as above; *e_g_* ∈ R^50^ is a vector of 50 single expression features, and *M_a_, M_b_*∈ R^1019^ are vectors of pseudobulk ChromVAR z-scores for singles A and B.

The model first encodes each peak in 256 dimensions (peak encoder) via a two-layer MLP with LayerNorm, dropout of 10%, and SiLU activation. Sinusoidal positional encoding is applied to the compressed peak representation to allow the model access to TSS distance information. Peak representations are then refined to account for local chromatin context via two TransformerEncoder blocks,^59^ consisting of pre-norm multi-head self-attention across peaks (4 heads) with a feed-forward network of hidden dimension 1024 and GELU activation. Single expression is incorporated via a FiLM layer,^87, 88^ in which a two-layer MLP with hidden dimension 512 maps the 50-element expression vector to 256-dimensional scale and shift parameters that further update the peak embeddings. A shared linear encoder is applied to ChromVAR z-scores for singles A and B, projecting each to 512 dimensions. We then apply the permutation-invariant concatenation operation as above to produce one “pair representation” for double AB as the 1024-dimensional vector [*A* + *B, A* × *B*]. This vector is updated via an MLP that projects to 256 dimensions via two fully connected layers of size 512 and 256, LayerNorm, SiLU activation, and dropout of 10%. The updated pair representation finally queries the refined peak representations via multi-head cross-attention (InteractionHead), and the resulting pooled representation is passed through an MLP with LayerNorm, hidden dimension 64, SiLU activation, and dropout of 10% to predict the final scalar residual t-statistic for double AB in gene *g*.

The model was implemented in PyTorch and trained to minimize a mean squared error loss between observed and predicted residual t-statistics, using the AdamW optimizer with weight decay 1e-6 and OneCycleLR scheduler with maximum learning rate 1e-5 for the peak encoder and PeakBlock, and 1e-4 for all other parameters. The batch size for training was 1,024 individual (gene, double) examples; training takes about 2 hours on 2 Nvidia A40 GPUs. For each of 12 cross-validation folds (split by chromosome as above), the model was trained for 20 epochs, and the checkpoint with the highest Pearson correlation between observed and predicted residuals on the validation set was used for inference. During training, we added 50 examples for each gene where the “double” consisted of ChromVAR information for the same single perturbation twice (e.g., A paired with A), setting the target residuals for these examples to 0. These examples were excluded from inference and performance calculation on the test set. For modality-specific ablation (Fig. 6D, Fig. S9D), each modality (peak matrix, single expression vectors, and ChromVAR z-scores) was replaced with the feature-wise training-set mean during inference. For evaluation of model performance at program level (Fig. 6C), program residuals for each pair were calculated as a weighted average of observed or predicted gene-level residual t-statistics using each gene’s program weight.

### Attribution of salient regions and chromatin features

To identify recurrent chromatin features driving the model’s predictions, we built a two-stage attribution pipeline to first nominate relevant chromatin regions for each gene, then quantify the importance of each motif and accessibility feature within the relevant regions. We first subset the gene universe to only consider program genes accurately predicted by the model (correlation of observed and predicted residuals across all doubles of at least 0.25) and for which more variance in residuals was explained by observed vs. ablated *cis*-chromatin information (*R*^2^ of predictions greater than that of peak-ablated predictions). We then ran the following pipeline for each gene, using the model checkpoint from the cross-validation fold in which its chromosome was in the test set. For each (gene, double) prediction, we first nominated 32 of the 128 peaks nearest the gene’s TSS as salient chromatin regions using the peak Jacobian score introduced by Fu et al.^60^ Briefly, for each peak, the gradient of the model’s output with respect to the peak encoder’s final layer (ideally less noisy than raw peak inputs) was calculated, and the L_2_ norm of this vector was taken to produce a scalar quantifying the peak’s importance. The top 32 peaks by this metric were selected for downstream attribution. IntegratedGradients attribution^89^ (multiply_by_inputs = False, n_steps = 10) implemented in the captum library was then applied to each of these peaks’ input features, holding expression and ChromVAR scores fixed, using the training set mean as the baseline. The final attribution matrix of shape 1,225 pairs × 464 peak features for each gene was then calculated by averaging the IntegratedGradients outputs across each of the top peaks and repeating the procedure for each pair.

To produce program-level attributions, we first concatenated each gene’s attribution matrix and L1-normalized across rows (to mitigate genes with large residual predictions from dominating the output), then z-normalized across columns (to align attribution scales for all motif and accessibility features). For each program, we aggregated these normalized attributions via a weighted sum using each gene’s sparse PCA program weight. Constitutive and conditional regulators (Fig. 7C) were nominated via the mean attribution of each feature across all pairs or the cosine similarity of each feature’s attribution score and the observed program residual for each pair. We finally validated program attributions by comparing top regulators to single-TF activation phenotypes in our previous work. Starting with the pre-processed RPE-1 TFome-wide CRISPRa expression matrix^90^, we pseudobulked by sgRNA and filtered to sgRNAs with observed on-target activation and no “bad seed” protospacer motifs. We scored the remaining subpopulations grouped by sgRNA on each of our programs as the weighted average of z-normalized expression using each gene’s program weight. For each program, we calculated the mean attribution score across all pairs for each motif cluster and accessibility feature, then mapped these model features to activated TFs to compare attribution scores to observed phenotypes (Fig. 7D). Accessibility features, which unambiguously implicate a single TF regulator, were mapped directly to that TF. If multiple sgRNAs against the TF were scored, the highest score was assigned if the feature’s attribution score was positive, else the lowest score if the attribution was negative. Motif cluster features were scored similarly, but all TFs whose motifs were in the cluster were considered for scoring in clusters containing more than one motif.

### TF triples screen data processing and residual calculation

RNA-seq data processing was performed via Plasmidsaurus’s in-house pipeline. Quality of raw fastq files was assessed using FastQC v0.12.1. Reads were then quality-filtered using fastp v0.24.0 with poly-X tail trimming, 3’ quality-based tail trimming, a minimum Phred quality score of 15, and a minimum length requirement of 50 bp. Quality-filtered reads were aligned to the reference genome using STAR aligner v2.7.11 with non-canonical splice junction removal and output of unmapped reads, followed by coordinate sorting using samtools v1.22.1. PCR and optical duplicates were removed using UMI-based deduplication with UMIcollapse v1.1.0. Alignment quality metrics, strand specificity, and read distribution across genomic features were assessed using RSeQC v5.0.4 and Qualimap v2.3, with results aggregated into a comprehensive quality control report using MultiQC v1.32. Gene-level expression quantification was performed using featureCounts (subread package v2.1.1) with strand-specific counting, multi-mapping read fractional assignment, exons and 3’ UTR as the feature identifiers, and grouped by gene_id. Final gene counts were annotated with gene biotype and other metadata extracted from the reference GTF file. Log fold changes for each condition with respect to control cells (single NTC guide) were calculated with edgeR v4.0.16 using standard practice including filtering for low-expressed genes with edgeR::filterByExpr with default values.

Starting with the edgeR-estimated LFC values, we calculated program residuals by first scoring the three single TFs, the FOSB_SOX18 double, and the three triple conditions on the expression of program genes, calculating scalar program scores *s* as the dot product of feature weights and expression values in each program. The limited number of conditions in the triples screen prevented us from fitting a global null expectation, so we empirically scaled the scores of the double and single conditions by √2 such that the L2 norm of the expectation’s coefficients is 1. This produces a distribution of program residuals *r* with mean and median close to 0, correcting for saturation effects. Program residuals were then calculated by subtracting the sum of scaled scores in double and single conditions from the triple’s score.

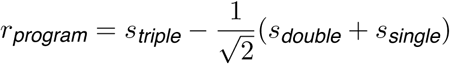

Gene-level residuals in log fold change space did not require a similar saturation correction, and a purely additive expectation given the LFC values *x* produced a distribution with mean and median close to 0.

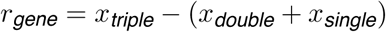

### Curation of AMD Genetic Association Studies

All GWAS summary statistics and fine-mapping were based on the European ancestry meta-analysis performed by Gorman et al.^50^, which is the most comprehensive GWAS meta-analysis and fine-mapping of AMD to date. This meta-analysis had 57,290 cases and 324,430 controls, and included individuals from the MVP, IAMGDC, GERA, UK Biobank, and Genentech GA and CNV cohorts. Summary statistics were downloaded from dbGaP study accession phs001672 using the SRA toolkit by taking the “AMD.EUR_META.NatGen2024.txt.gz” file. Fine-mapping results were obtained from the Gorman et al. study.^50^ Fine-mapping identifies credible sets, which are groups of variants that collectively contain a likely casual variant. Briefly, fine-mapping was performed in Gorman et al. using the FINEMAP^91^ method on genome-wide significant loci. Variants in both the summary statistics and fine-mapping results were filtered to common and low frequency variants (minor allele count ≥ 5) from the 1000 Genomes Project.^53^

For the genetic epistasis analysis described below (see **Genetic epistasis informed by Multiome Perturb-seq**), phenotype information for AMD was obtained from data fields 30926 (left eye) and 30907 (right eye). This AMD phenotype was derived from manual grading of retinal optical coherence tomography scans performed across 68,514 participants.^92^ This data only became available on the UK Biobank in the November 2025 release. Notably, this phenotype has 10,457 cases and 10,492 cases for the left and right eye, respectively. This expands on other data fields containing self-reported AMD phenotypes (e.g. data field 5923 has 4,757 cases and data field 6148 has 7,815 cases). We reasoned that this expanded number of AMD cases could provide a more robust phenotype for nominating genetic epistasis hypotheses. All genetic epistasis analyses were performed on the UK Biobank Research Analysis Platform under application 16549.

### Calling peak sets across perturbations

We applied scE2G^54^ to call peaks and create peak-gene maps for each perturbed cell subpopulation. Briefly, scE2G is a logistic regression classifier model that uses information about chromatin accessibility, gene expression, and genomic features as input features, and is trained to predict CRISPR-validated enhancer-gene interactions in a leave-one-chromosome out fashion. Within a single-cell dataset, scE2G is designed to create a peak-gene map within a single cell type or cluster. Here, we treat different perturbations as clusters and run scE2G on each unique perturbation (e.g. one peak set and peak-gene map for cells with *PRDM16* activated, one peak-gene map for cells with *OTX2* activated, etc.). Following the guidance of scE2G, we only generated peak-gene maps for perturbations with more than 100 cells, where scE2G shows robust performance. This allowed us to create peak-gene maps for 55 of the 120 perturbations. All peak-gene maps were generated using the scE2G^Multiome^ model.

Peak sets were called on each perturbed cell subpopulation using scE2G, which uses a MACS2-based^93^ peak calling approach to define candidate regulatory elements (more details in the scE2G paper^54^). There were 55 peak sets corresponding to 55 perturbations with greater than 100 cells. “All peaks” were defined as the union of all peak sets. “Control peaks” were defined as the peaks in control cells. “Perturbed peaks” (or perturbation-specific regions) were defined as peaks in cell populations receiving a gene perturbation but removing peaks that have any overlap with control peaks. The respective “linked” peak sets were derived the same way, but subsetting to peaks linked to genes by scE2G.

To avoid linkage disequilibrium-related inflation when assessing the number of AMD loci revealed by each perturbation, we defined perturbation-specific credible sets. To assign credible sets to perturbations, we first marked all credible sets where a variant with PIP *>* 0.1 overlaps a control peak as a “control credible set.” Then, we defined perturbation-specific credible sets as all credible sets where a variant with PIP *>* 0.1 overlaps a peak in any perturbed cell population and the credible set is not in the control credible sets. We assigned variants to credible sets using the credible set assignments from the FINEMAP^91^ run with the maximum number of causal signals allowed in a locus where at least one variant in the locus was assigned to a credible set.

### Perturb-PoPS model for polygenic-monogenic convergence

We adapted the PoPS framework^57^ for our Perturb-seq features and call the predicted scores from this approach “Perturb-PoPS”. Perturb-PoPS uses a combination of gene-level GWAS associations with gene-level features from a Perturb-seq experiment to nominate disease genes that are co-regulated with polygenic GWAS genes in the Perturb-seq experiment. Here, we apply Perturb-PoPS both across the entire set of TF perturbations (both pairs and single TFs) in our dual guide experiment, but also to subsets of TF pairs. Specifically, we are interested in assessing whether there is co-regulation between polygenic and monogenic genes within Perturb-seq data.

### Feature curation

Analogous to the PoPS framework^57^, we created gene-level features for Perturb-PoPS. While the original PoPS derived these features from gene expression datasets, gene pathways, and protein-protein interactions, here we derive these features directly from our dual guide Perturb-seq experiment. We first take the matrix of z-normalized gene expression measurements per cell (see **Dual guide Perturb-seq: data processing and quality control**). We then regress out cell cycle phase, library size, and mitochondrial read percentage (using `sc.pp.regress_out()`). Finally, we pseudobulk these adjusted expression values per perturbation such that we have a gene-level feature for each single and double perturbation in our dual-guide experiment. Notably, gene-level features from our Perturb-seq experiment could only be generated for those passing our expression thresholds. Therefore, all analyses involving Perturb-PoPS are subsetted to genes with both MAGMA^56^ scores and passing the expression threshold in our dual guide Perturb-seq dataset.

### Training Perturb-PoPS model

We first define the subset of TF perturbations that we are interested in running Perturb-PoPS on:

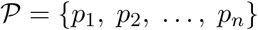

Here, *p*_1_ is one perturbation taken from our dual guide Perturb-seq experiment and can be from a single TF perturbation or a TF pair perturbation. We then perform a feature selection step where each feature is individually regressed against the AMD MAGMA scores. For a given perturbation *p* in the larger perturbation set P, the model is:

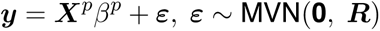

Here, ***y*** is a vector of the per-gene MAGMA scores, ***X****^p^* is a vector of the per-gene z-normalized expression measurements for perturbation *p* in the dual guide Perturb-seq experiment, *β^p^* is the coefficient estimated by fitting the model, ***ε*** is an error term, and ***R*** is a covariance matrix that accounts for LD between nearby genes obtained from MAGMA. This model is fit using generalized least squares (GLS). We obtain a *p*-value for the alternate hypothesis that *β^p^* ≠ 0. We retain all perturbations that pass a nominal significance threshold (*p <* 0.05) for the joint PoPS model (second step).

We next fit the joint PoPS model where all selected features from the previous step are jointly regressed against the AMD MAGMA scores. Here, we can also subset the perturbations that are included in the joint model to a perturbation subset P as we did in the analyses presented in Fig. 5C. The joint model is:

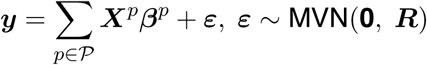

Here, P is the subset of perturbations from the dual guide Perturb-seq experiment to be included in Perturb-PoPS that also passed the feature selection step described above. This model was fit using GLS with L_2_ regularization. Importantly, the joint model was fit in a leave-one-chromosome-out (LOCO) fashion. This means that for a given chromosome, the joint model was fit on all the other chromosomes and then used to generate predictions for the given chromosome. This procedure yields a separate fitted joint model for each chromosome. Therefore, the Perturb-PoPS score for a given gene *g* can be represented as:

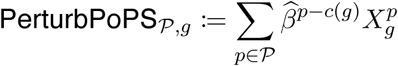

Here, PerturbPoPS*P,g* represents the Perturb-PoPS score for a perturbation subset P and gene *g*. *c* (*g*) represents the chromosome that gene *g* is on, such that *β^p−c^*^(^*^g^*^)^ represents the coefficient learned from the joint model for the perturbation from all other chromosomes. *X^p^* represents the z-normalized gene expression measurement from the dual guide Perturb-seq experiment for gene *g* in perturbation *p*. Prior to fitting the model, we standardize features.

For the Perturb-PoPS scores generated from singles features only, the perturbation subset P was restricted to single TF perturbation in the dual guide Perturb-seq experiment. For the doubles score, Pincluded both single TF and TF pair perturbations. For the analysis in Fig. 5C, P was restricted to TF pair perturbations within the marked cluster. Clusters were generated using Ward clustering with a Euclidean distance metric.

For comparisons with the original PoPS method, we recomputed the AMD PoPS score using the original feature set, but subsetting to the genes expressed in the dual guide Perturb-seq experiment. This ensures that both Perturb-PoPS and the original PoPS features can learn disease-relevant signal across the same gene sets. However, this introduces a limitation with Perturb-PoPS where scores are generated on a smaller gene set than the original PoPS approach.

To generate a PoPS score using features derived from control RPE-1 cells, we adapted the PoPS feature generation pipeline (available at https://github.com/FinucaneLab/gene_features) and applied it to the raw counts matrix from the RPE dual guide experiment subsetted to the control cells. We first generated the features and then subsetted to the genes matching our expression threshold for the dual guide experiment. Features include principal components and independent components (both within clusters and across all cells), average scaled expression per cluster, and differential expression in clusters. More details can be found in Weeks et al.^57^

### Enrichment of monogenic genes for Perturb-PoPS

For computing the recall of monogenic genes (Mendelian retinal disease genes), we only considered monogenic genes that we generated Perturb-PoPS, PoPS on RPE-1 controls, and original PoPS (subsetting genes) for. We then considered recall of monogenic genes both in the top 500 and top 1,000 PoPS genes. We computed enrichment of monogenic genes as the following:

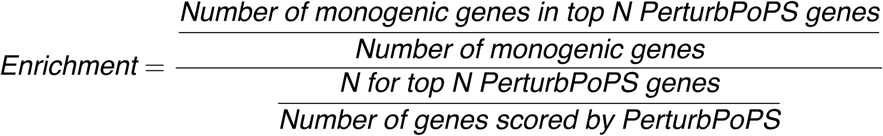

Recall was computed as the following:

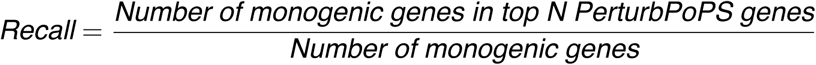

We then computed jackknife enrichments and recalls using a leave-one-chromosome out approach and then computed a standard error for the enrichment and recall, and computed p-values using a one-sided hypergeometric test.

### Curation of relevant disease gene sets for AMD

MAGMA scores^56^ for each gene are computed from summary statistics by taking the mean chi-squared test statistic for SNPs within a specified window of the gene body as a test statistic. For each gene, a Z-score (the MAGMA score) is computed by creating a null distribution of test statistics that accounts for within-gene LD and comparing the observed test statistic to the null distribution. We applied MAGMA directly on the AMD meta-analysis summary statistics downloaded from dbGaP (see **Curation of AMD Genetic Association Studies**). We then filtered genes on the X chromosome. These gene-level Z-scores and gene-gene correlation matrix from MAGMA also served as downstream inputs to PoPS. For determining MAGMA genes in downstream analyses, we consider all genes with a ZSTAT *>* 3.

PoPS^57^ scores are computed by training a linear model to predict MAGMA Z-scores using features derived from a combination of gene expression datasets (both single-cell and bulk), protein-protein interactions, and gene pathways. The linear predictor from this linear model is then used as the PoPS score to additionally prioritize potential disease genes that are in the same functional pathways as GWAS genes. We generated PoPS scores for AMD using our previously computed MAGMA scores. For the other 113 complex traits that we compared to AMD, we downloaded the PoPS results generated by Weeks et al.^57^ from the Finucane lab website (https://www.finucanelab.org/data).

To identify a relevant monogenic gene set for AMD, we used the Mendelian genes curated by Freund et al.^58^ Briefly, Mendelian genes were defined as fully penetrant OMIM genes with a known molecular basis. In their study, they then matched complex traits to matched Mendelian phenotypes. Here, we take all the Mendelian genes that were phenotypically matched to AMD, which we call Mendelian retinal disorder (MRD) genes.

### Pooled combinatorial interaction scores

We defined the pooled combinatorial interaction (PCI) score for each gene as the L_2_ norm of the residuals for that gene taken across all perturbations (see **Calculation of gene-level residuals**). For each our three disease gene sets (PoPS, MAGMA, and Mendelian), we compared the PCI scores of the disease genes to those that were not, using the respective definitions of disease genes. Within each gene group, we took the mean and standard error of the PCI score. P-values for whether disease genes showed elevated PCI scores were estimated with a one-sided t-test. For determining whether individual TF pairs showed elevated residuals at disease gene sets, we replaced the PCI scores with the residuals for the individual TF pairs.

PoPS scores for 113 other complex traits were obtained directly from the Finucane lab website (see **Generation of PoPS gene-level disease prioritization scores for AMD**). For each trait, we took the top PoPS genes in the 1^st^, 5^th^, 10^th^, 15^th^, 25^th^, and 50^th^ percentiles. We then took the mean PCI score for the PoPS genes within each of these percentiles and the mean PCI score for all other genes and took the difference of these two means for each percentile. We then took the average of these differences across all the percentiles to get a “mean difference” metric for each trait, representing the elevation of PCI scores in the top PoPS genes relative to other genes for that trait.

### Genetic epistasis informed by Multiome Perturb-seq

Previous studies in genetic epistasis testing have found that genetic epistasis in a variant pair can be explained by weak LD of one of the variants with a third variant that has a strong additive effect on the phenotype causing inflation of test statistics, particularly when testing two variants in proximity with each other, but also when testing two unlinked variants on different chromosomes.^94, 95^ In our genetic epistasis tests here, we focus on variants on different chromosomes, exclusively focus on variants in fine-mapped credible sets, and are using a complex trait phenotype. However, we caution the reader that genetic epistasis tests are complex and can be susceptible to residual LD-related artifacts, potentially leading to false positives, as demonstrated previously.^94, 95^

To nominate a larger set of variants for genetic epistasis testing, we first performed a genome-wide fine-mapping using SuSIE^96, 97^ on our AMD summary statistics (see **Curation of AMD genetic association studies**). We applied SuSIE on 1,361 approximately independent autosomal LD blocks from the Alzheimer’s Disease Sequencing Project (ADSP)^98^. Each LD block is extended by 50 kb on both sides to define a fine-mapping region and reduce boundary effects for signals near block edges. This identified 3,124 variants within credible sets.

We next used functional information from our Multiome Perturb-seq experiment to restrict the variant pairs that we test for genetic epistasis. We took all peaks from SnapATAC2^79^ (see **Multiome Perturb-seq: data processing and quality control**) that were differentially accessible in response to at least 1 guide RNA (*n* = 333,681 peaks, 561,867 guide-peak pairs). We then intersected variants in SuSIE credible sets with the differentially accessible peaks (*n* = 266 variant-peak pairs). However, we deduplicated cases where multiple variants from the same credible set overlap a peak by keeping the peak overlapping the variant with the highest PIP in the credible set, resulting in 58 variant-peak pairs. Merging these variant-peak pairs with the guides that induce differential accessibility of peaks resulted in 107 guide-variant-peak sets. We then took pairwise combinations of variants that are differentially accessible in response to the same guide RNA, resulting in 260 guide-variant-variant sets. From here, 227 unique variant-variant pairs nominated for further genetic epistasis testing.

To test for genetic interactions, we used the UK Biobank cohort.^99^ Variant genotypes were retrieved using plink2.^100, 101^ Individual-level AMD phenotypes were retrieved from the UK Biobank using field IDs 30926 (left eye AMD, 10,457 cases) and 30907 (right eye AMD, 10,492 cases) (see **Curation of AMD genetic association studies** for more details). In our analysis, an individual was marked as a case for AMD if they had AMD in either eye or marked as a control otherwise. Individuals with genotypes that were not 0, 1, or 2 for both variants were removed.

We further filtered individuals included in our genetic epistasis testing to individuals of white British ancestry who are age 65 and older (mirroring the Gorman et al.^50^ procedure). We also removed individuals who did not have a value for standing height. Furthermore, we binarized genotypes for any variant with a MAF *<* 0.05.

We considered 3 different allele codings for each variant - dominant, recessive and additive. We illustrate our genetic epistasis test for the recessive-recessive scenario that generated a nominally significant epistatic effect we report in Fig. S6. In this scenario, individuals with two copies of an allele for a particular variant are marked 1 and all other individuals are marked 0. We first analyze the proportion of cases in each of the 4 possible genotype combinations (0/0, 0/1, 1/0, and 1/1) to see if there are differences in the proportions. Error bars show the 95% confidence interval based on a Wilson score interval.

We next performed a stratified genetic epistasis test to get a p-value for the difference in proportions between two genotypes conditional on another variant’s genotype. For each variant pair, we fit two models for each variant, stratifying conditional on the other variant (i.e., we test for the effect of variant 2 when the genotype of variant 1 is 1, and then we test for the effect of variant 2 when the genotype of variant 1 is 0). We then repeat this for the other variant in the variant pair. This can be represented as follows with 4 models:

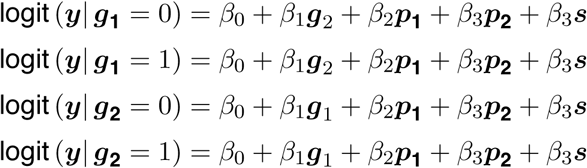

Here, ***y*** represents the AMD phenotype across individuals, ***g*_1_**and ***g*_2_** represent the genotype for either variant 1 or 2, *β*_0_ represents an intercept term, *β*_1_ is the coefficient for the variant effect on AMD (variant 2 or 1), ***p*_1_** is the vector of genotype principal component (PC) 1 across individuals, *β*_2_ is the coefficient for PC 1, *β*_3_ is the coefficient for PC2, ***p*_2_** is a vector of genotype PC 2 across individuals, ***s*** is a vector of sex across individuals, and *β*_3_ is the coefficient for sex.

Genotypes and covariates were standardized prior to fitting the models. We included the first two genotype PCs and sex as covariates, mirroring the procedure from Gorman et al. *P* -values were obtained for the *β*_1_ coefficient using a Wald test from the `glm()` function in R. Based on the QQ-plot, we did not observe inflation of our *p*-values. We also removed cases where one of the variants is in the HLA region (chromosome 6:25.5-34.0 Mb)^102^ and if both variants are on the same chromosome. We view this suggestive hit as an exploratory proof of principle as purely hypothesis-generating rather than establishing evidence of genetic epistasis.

### Enrichment and recall of AMD fine-mapped variants

We analyzed the 301 fine-mapped AMD variants from Gorman et al.^50^ (see **Curation of AMD Genetic Association Studies** for more details). For a given peak set, we defined enrichment as:

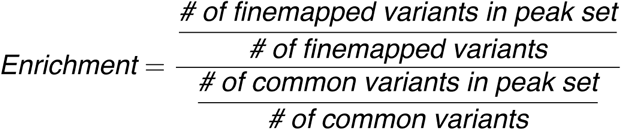

Here, common variants were defined as common and low-frequency (minor allele count ≥ 5) variants from the 1000 Genomes Project.^53^

Recall was defined as:

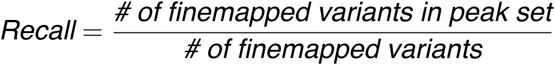

Standard errors for the enrichment and recall were estimated through computing a sample standard deviation across 100 bootstrap enrichments or recalls by resampling chromosomes. P-values were computed using a Wald test by first estimating the standard error across 100 bootstraps and then computing a z-score using the formula:

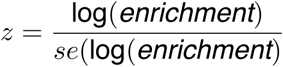

Then, two-tailed p-values were obtained using the CDF of the normal distribution.

### Heritability enrichment analyses

Heritability enrichment analyses were performed using the Stratified Linkage Disequilibrium Score Regression (S-LDSC) method (https://github.com/bulik/LDSC).^48^ First, each peak set (either cell type peaks from Wang et al. or RPE-1 bulk ATAC-seq peaks) was converted to a variant-level annotation (a binary 0/1 indicating whether a SNP overlaps a peak) using the 1000 Genomes Project European phase 3 variants covering 9,991,229 variants with minor allele count ≥ 5.^53^ Partitioned LD scores were then computed for each peak set annotation using the 1000 Genomes Project European samples, an LD window of 1 centimorgan, and only printing LD scores for HapMap3 SNPs.^103^ Prior to LD score regression, p-values of 0 in the summary statistics had to be replaced with 1e-300 so that the S-LDSC could be run. Stratified LD score regression was then performed separately on each peak set, conditioning on the baseline model from Finucane et al.^48^ containing 53 annotations covering coding and regulatory annotations. Here, the baseline model ensures more unbiased estimates of heritability enrichment. Allele frequencies were taken from the 1000 Genomes Project Phase 3 European samples, and LD scores from HapMap3 SNPs excluding the HLA region were used as weights in the regression.

Briefly, heritability enrichment is defined as:

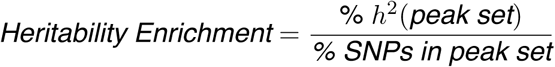

where % h^2^(peak set) is the proportion of AMD heritability captured by SNPs overlapping peaks in the peak set.

### Enrichment of disease genes in gene programs

For each program, we first took the list of all genes within the gene program. As a background, we considered all genes that could possibly be included in the program creation (see **Generation of gene programs by sparse PCA**). For PoPS, we additionally subsetted genes to those for which we generated a PoP score. To determine the PoPS genes here, we took the top 500 PoPS genes for AMD, and then subsetted to those included in the program creation for the dual guide experiment. We then defined *a* = *length*(*program and disease genes*), *b* = *length* (*program genes*)−*a*, *c* = *length* (*disease genes*)−*a*, and *d* = *length* (*background genes*)−*a*−*b* −*c*. We then ran a one-sided Fisher’s exact test to test whether the program was enriched for disease genes, and computed the odds ratio using the following formula:

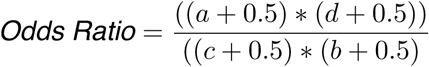

We used the Haldane-Anscombe correction here to keep odds ratios finite in case *b* or *c* is 0.

### Bulk ATAC-seq peaks in RPE-1 cells

Bulk ATAC-seq in RPE-1 cells from Southard*, Ardy* et al. was performed in 3 replicates. We applied ’macs3 callpeak -f bam -g hs --nomodel --shift -100 --extsize 200 -q 0.05’ to call peaks across all 3 replicates jointly using MACS3.^104^ The bulk RPE-1 WT cell ATAC-seq peaks from the 4D Nucleome Project (“Bulk peaks (4DN)”) were downloaded from the 4DN Data Portal (4DNES48BA2C8). This data was generated in the Bing Ren lab.^51, 52^

