## Supplementary material for "Engineered regulatory states resolve context-dependent genetics in macular degeneration": Document S1

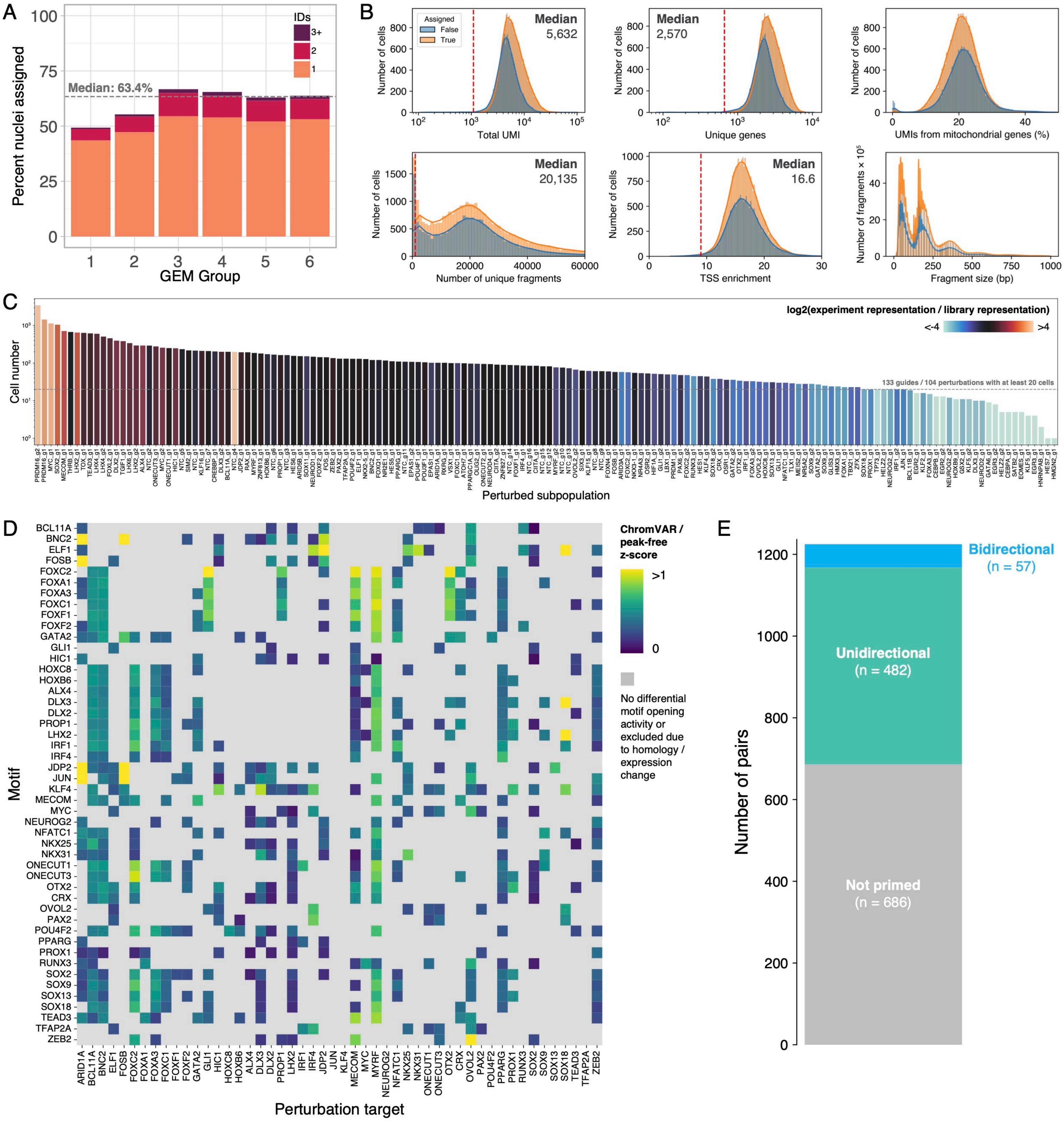

A
Percent nuclei assigned
Median: 63.4%
IDs
3+
2
1
GEM Group
B
Assigned
False
True
Median
5,632
Number of cells
Total UMI
Median
2,570
Unique genes
UMIs from mitochondrial genes (%)
Median
20,135
Number of unique fragments
Median
16.6
TSS enrichment
Number of fragments × 10⁵
Fragment size (bp)
C
log2(experiment representation / library representation)
<-4
>4
Cell number
133 guides / 104 perturbations with at least 20 cells
Perturbed subpopulation
D
ChromVAR / peak-free z-score
>1
0
No differential motif opening activity or excluded due to homology / expression change
Motif
BCL11A
BNC2
ELF1
FOSB
FOXC2
FOXA1
FOXA3
FOXC1
FOXF1
FOXF2
GATA2
GLI1
HIC1
HOXC8
HOXB6
ALX4
DLX3
DLX2
PROP1
LHX2
IRF1
IRF4
JDP2
JUN
KLF4
MECOM
MYC
NEUROG2
NFATC1
NKX25
NKX31
ONECUT1
ONECUT3
OTX2
CRX
OVOL2
PAX2
POU4F2
PPARG
PROX1
RUNX3
SOX2
SOX9
SOX13
SOX18
TEAD3
TFAP2A
ZEB2
ARID1A
BCL11A
BNC2
ELF1
FOSB
FOXC2
FOXA1
FOXA3
FOXC1
FOXF1
FOXF2
GATA2
GLI1
HIC1
HOXC8
HOXB6
ALX4
DLX3
DLX2
PROP1
LHX2
IRF1
IRF4
JDP2
JUN
KLF4
MECOM
MYC
MYRF
NEUROG2
NFATC1
NKX25
NKX31
ONECUT1
ONECUT3
OTX2
CRX
OVOL2
PAX2
POU4F2
PPARG
PROX1
RUNX3
SOX2
SOX9
SOX13
SOX18
TEAD3
TFAP2A
ZEB2
Perturbation target
E
Number of pairs
Bidirectional
(n = 57)
Unidirectional
(n = 482)
Not primed
(n = 686)

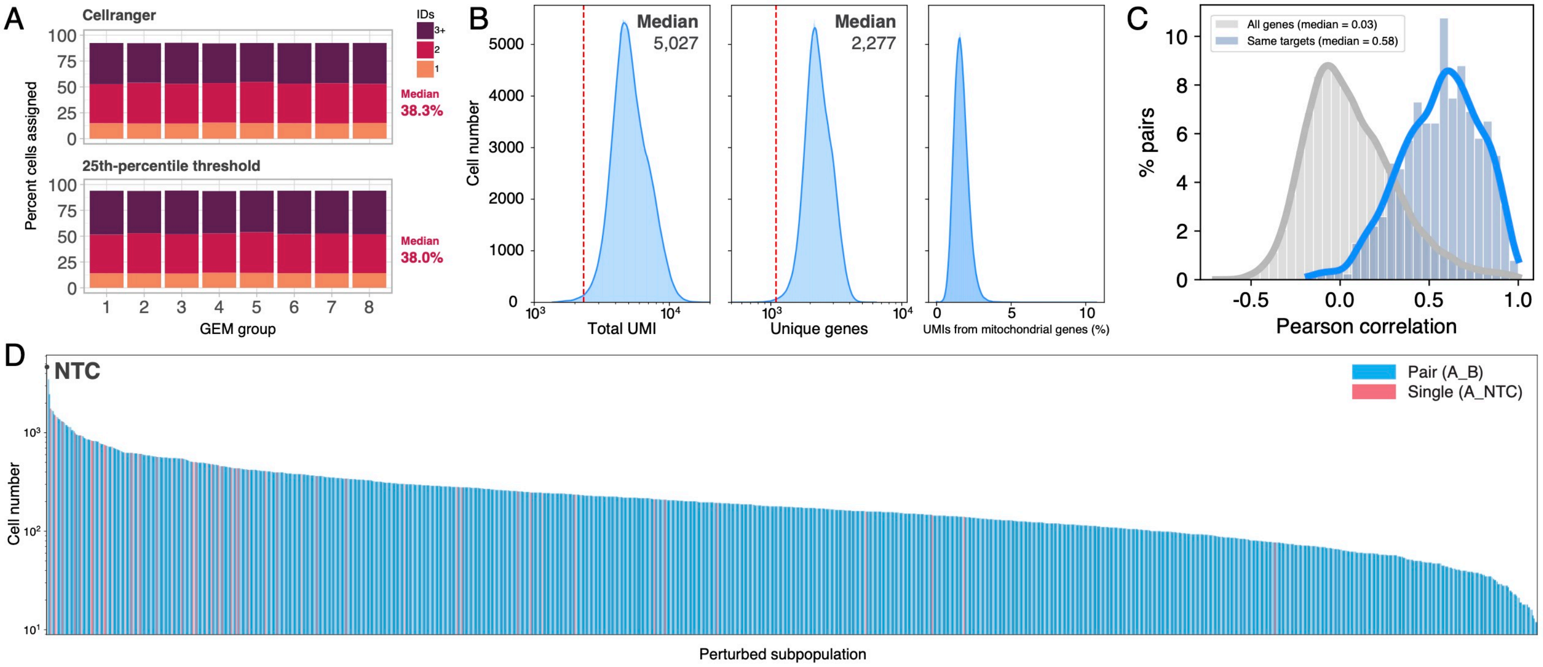

A
Cellranger
25th-percentile threshold
Percent cells assigned
GEM group
IDs
3+
2
1
Median
38.3%
Median
38.0%
B
Cell number
Median
5,027
Median
2,277
Total UMI
Unique genes
UMIs from mitochondrial genes (%)
C
All genes (median = 0.03)
Same targets (median = 0.58)
% pairs
Pearson correlation
D
NTC
Pair (A_B)
Single (A_NTC)
Cell number
Perturbed subpopulation

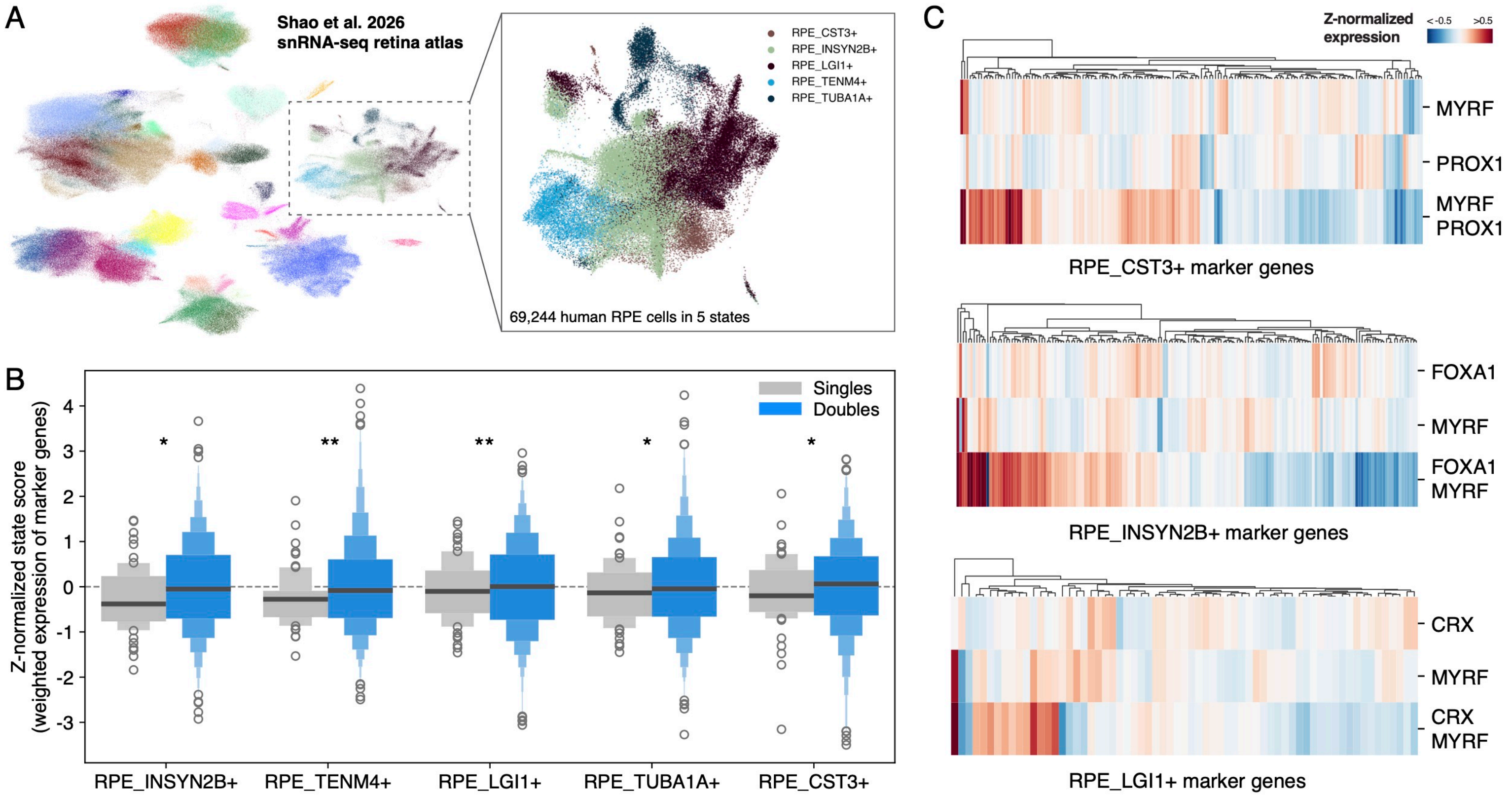
A
Shao et al. 2026
snRNA-seq retina atlas
RPE_CST3+
RPE_INSYN2B+
RPE_LGI1+
RPE_TENM4+
RPE_TUBA1A+
69,244 human RPE cells in 5 states
B
Z-normalized state score
(weighted expression of marker genes)
Singles
Doubles
RPE_INSYN2B+
RPE_TENM4+
RPE_LGI1+
RPE_TUBA1A+
RPE_CST3+
C
Z-normalized expression
< -0.5
>0.5
MYRF
PROX1
MYRF PROX1
RPE_CST3+ marker genes
FOXA1
MYRF
FOXA1 MYRF
RPE_INSYN2B+ marker genes
CRX
MYRF
CRX MYRF
RPE_LGI1+ marker genes

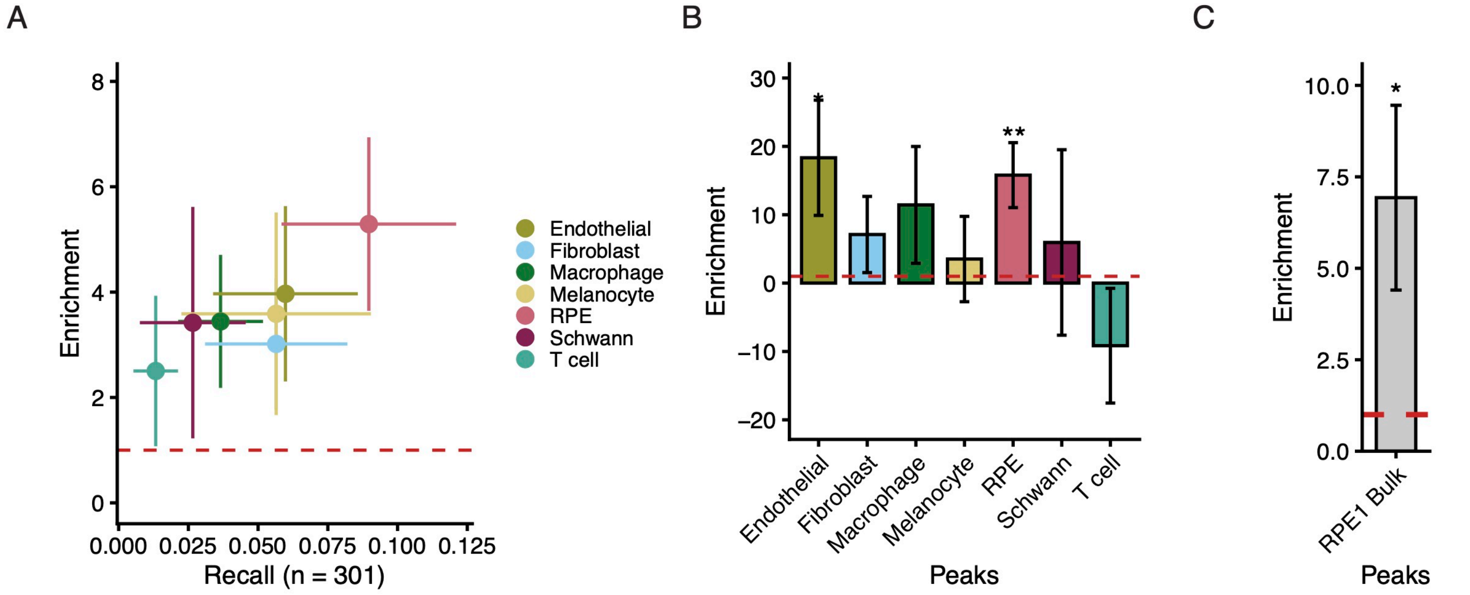
A
B
C
Enrichment
Recall (n = 301)
0.000
0.025
0.050
0.075
0.100
0.125
0
2
4
6
8
Endothelial
Fibroblast
Macrophage
Melanocyte
RPE
Schwann
T cell
30
20
10
0
−10
−20
*
**
Peaks
10.0
7.5
5.0
2.5
0.0
RPE1 Bulk

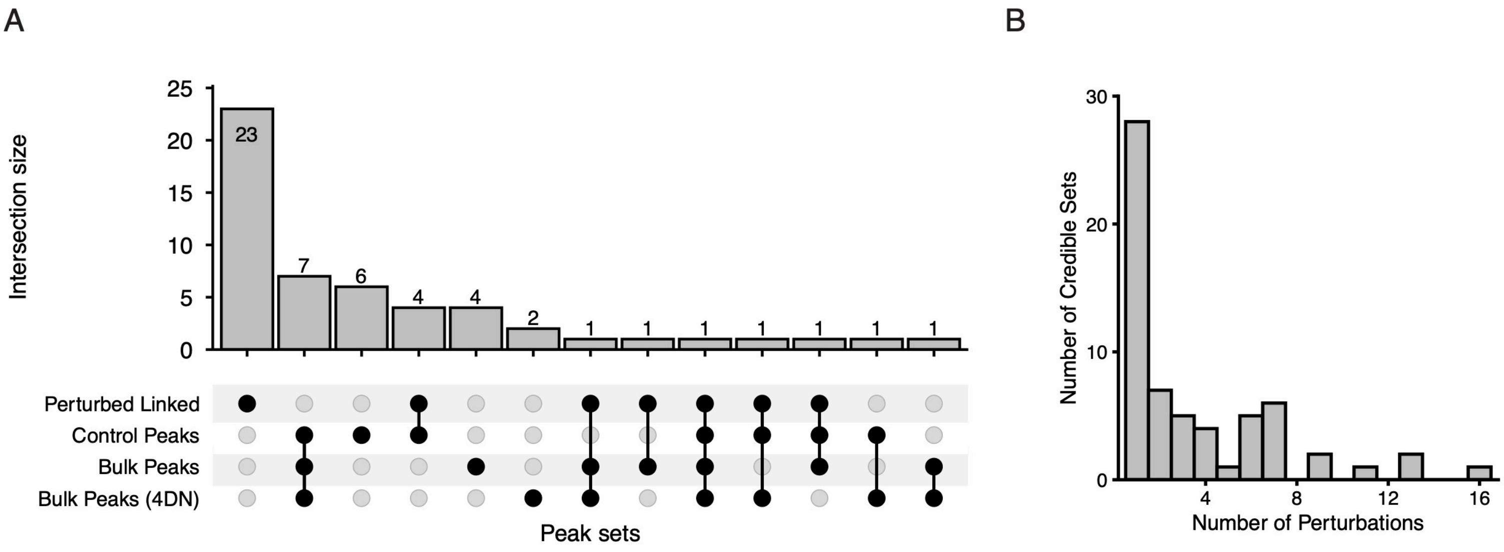
A
Intersection size
25
20
15
10
5
0
23
7
6
4
4
2
1
1
1
1
1
1
1
Perturbed Linked
Control Peaks
Bulk Peaks
Bulk Peaks (4DN)
Peak sets
B
Number of Credible Sets
30
20
10
0
4
8
12
16
Number of Perturbations

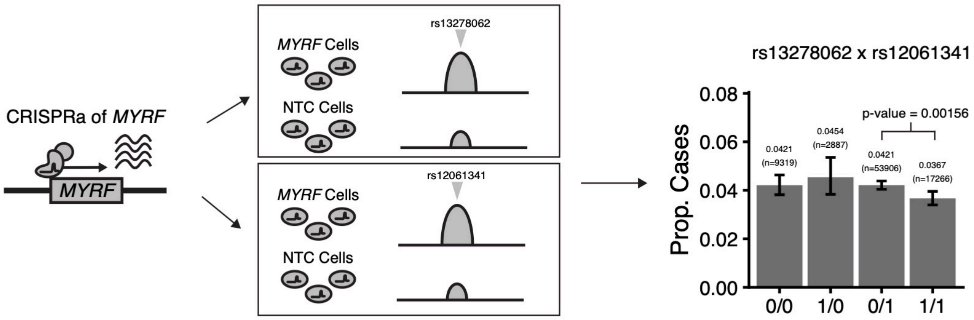
CRISPRa of MYRF
MYRF
MYRF Cells
NTC Cells
rs13278062
rs12061341
rs13278062 x rs12061341
Prop. Cases
p-value = 0.00156
0.0421
(n=9319)
0.0454
(n=2887)
0.0421
(n=53906)
0.0367
(n=17266)
0.00
0.02
0.04
0.06
0.08
0/0
1/0
0/1
1/1

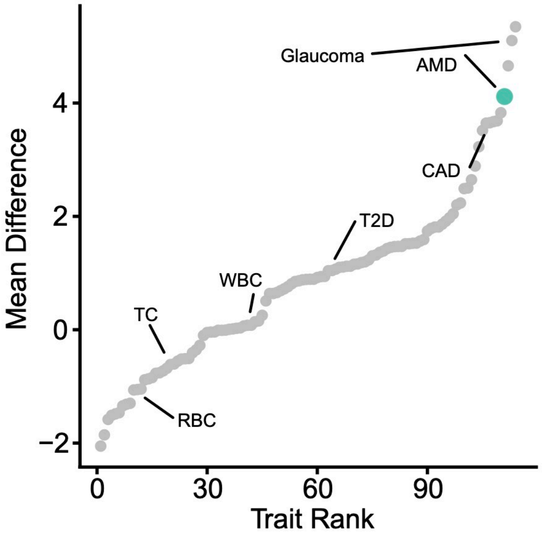

Glaucoma
AMD
CAD
T2D
WBC
TC
RBC
Mean Difference
Trait Rank
4
2
0
−2
0
30
60
90

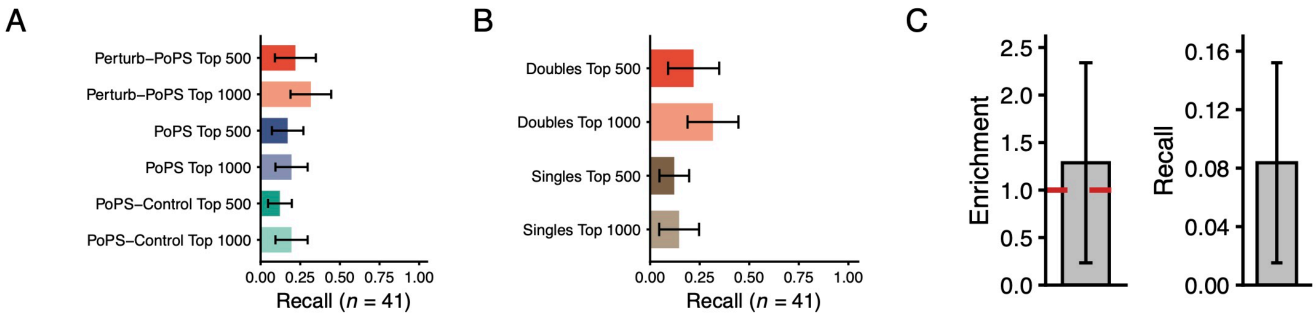

A
Perturb−PoPS Top 500
Perturb−PoPS Top 1000
PoPS Top 500
PoPS Top 1000
PoPS−Control Top 500
PoPS−Control Top 1000
0.00 0.25 0.50 0.75 1.00
Recall (n = 41)
B
Doubles Top 500
Doubles Top 1000
Singles Top 500
Singles Top 1000
0.00 0.25 0.50 0.75 1.00
Recall (n = 41)
C
Enrichment
0.0 0.5 1.0 1.5 2.0 2.5
Recall
0.00 0.04 0.08 0.12 0.16

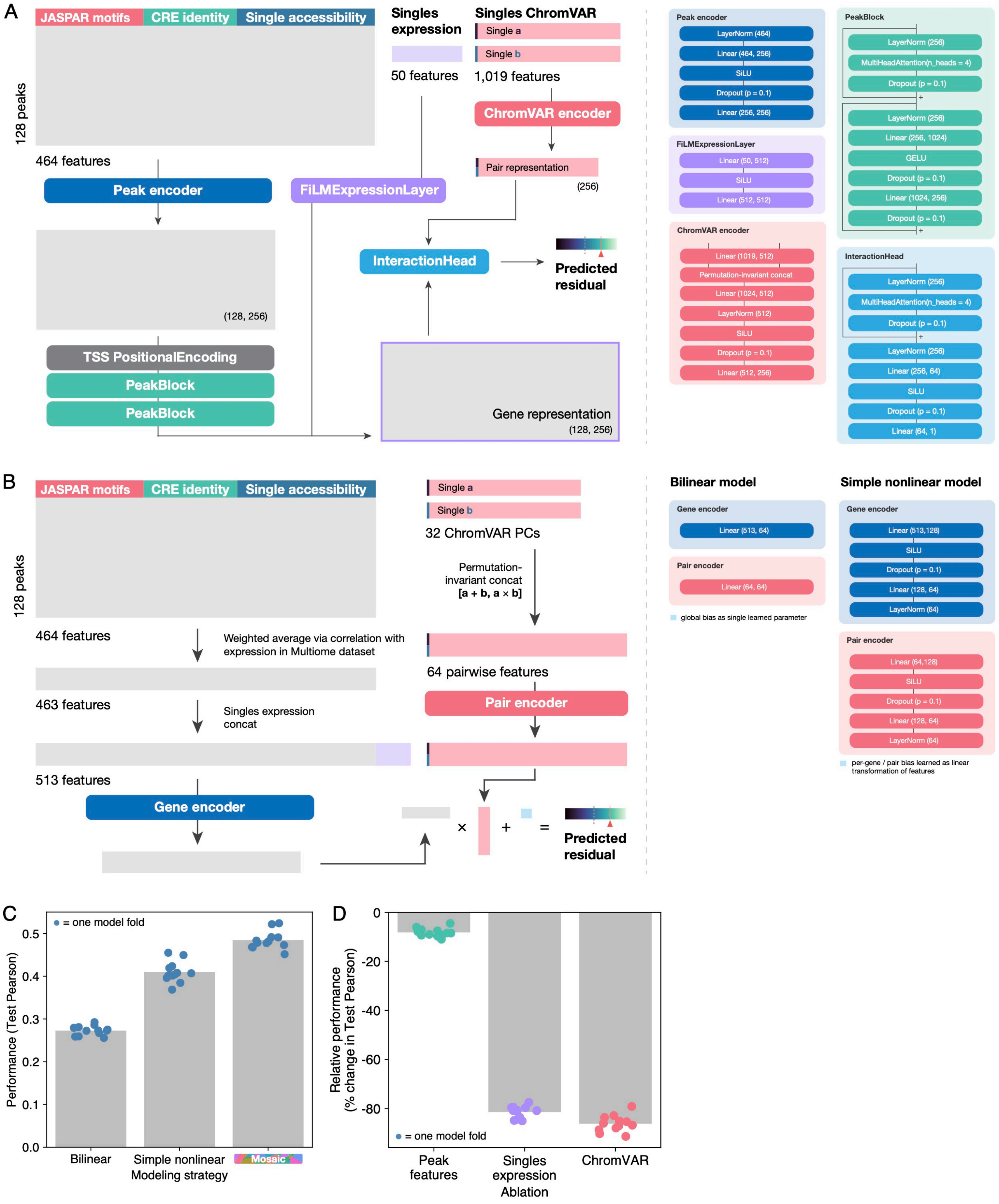
A
JASPAR motifs
CRE identity
Single accessibility
128 peaks
464 features
Singles expression
50 features
Singles ChromVAR
Single a
Single b
1,019 features
ChromVAR encoder
Pair representation
(256)
Peak encoder
FiLMExpressionLayer
InteractionHead
Predicted residual
(128, 256)
TSS PositionalEncoding
PeakBlock
PeakBlock
Gene representation
(128, 256)
Peak encoder
LayerNorm (464)
Linear (464, 256)
SiLU
Dropout (p = 0.1)
Linear (256, 256)
FiLMExpressionLayer
Linear (50, 512)
SiLU
Linear (512, 512)
ChromVAR encoder
Linear (1019, 512)
Permutation-invariant concat
Linear (1024, 512)
LayerNorm (512)
SiLU
Dropout (p = 0.1)
Linear (512, 256)
PeakBlock
LayerNorm (256)
MultiHeadAttention(n_heads = 4)
Dropout (p = 0.1)
LayerNorm (256)
Linear (256, 1024)
GELU
Dropout (p = 0.1)
Linear (1024, 256)
Dropout (p = 0.1)
InteractionHead
LayerNorm (256)
MultiHeadAttention(n_heads = 4)
Dropout (p = 0.1)
LayerNorm (256)
Linear (256, 64)
SiLU
Dropout (p = 0.1)
Linear (64, 1)
B
JASPAR motifs
CRE identity
Single accessibility
128 peaks
Single a
Single b
32 ChromVAR PCs
Permutation-invariant concat [a + b, a × b]
464 features
Weighted average via correlation with expression in Multiome dataset
463 features
Singles expression concat
64 pairwise features
Pair encoder
513 features
Gene encoder
×
+
=
Predicted residual
Bilinear model
Gene encoder
Linear (513, 64)
Pair encoder
Linear (64, 64)
global bias as single learned parameter
Simple nonlinear model
Gene encoder
Linear (513,128)
SiLU
Dropout (p = 0.1)
Linear (128, 64)
LayerNorm (64)
Pair encoder
Linear (64,128)
SiLU
Dropout (p = 0.1)
Linear (128, 64)
LayerNorm (64)
per-gene / pair bias learned as linear transformation of features
C
= one model fold
Performance (Test Pearson)
0.0
0.1
0.2
0.3
0.4
0.5
Bilinear
Simple nonlinear
Mosaic
Modeling strategy
D
Relative performance (% change in Test Pearson)
0
-20
-40
-60
-80
= one model fold
Peak features
Singles expression
ChromVAR
Ablation

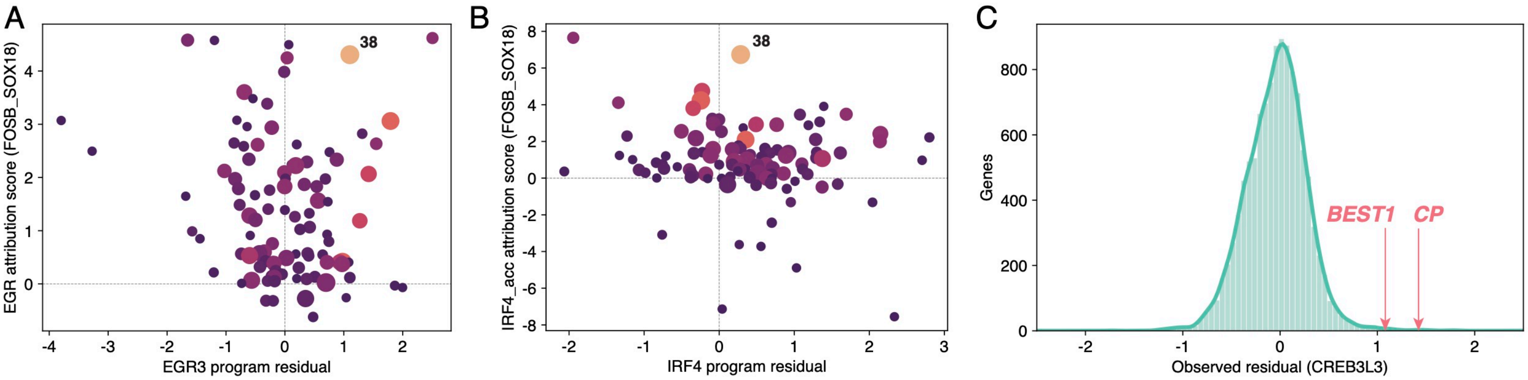
A
38
EGR attribution score (FOSB_SOX18)
EGR3 program residual
B
38
IRF4_acc attribution score (FOSB_SOX18)
IRF4 program residual
C
BEST1
CP
Genes
Observed residual (CREB3L3)

### SUPPLEMENTAL FIGURE LEGENDS

#### Figure S1: Quality control metrics related to the Multiome Perturb-seq single-TF screen

A. Percentage of single nuclei assigned to 1, 2, or 3+ sgRNA identities in each GEM group. Horizontal line shows median assignment rate across all GEM groups.
B. Gene expression (top) and ATAC (bottom) quality control metrics for single nuclei assigned or not assigned at least one sgRNA. Dashed lines indicate thresholds for filtering in final dataset used for analysis.
C. Number of cells maintained for analysis assigned to each sgRNA (25,129 total cells). Bar colors show change in sgRNA representation from initial plasmid library to experiment, a proxy for fitness effects.
D. Motif activity in primed TF pairs, as in Fig. 1H.
E. Number of non-primed, unidirectionally primed, and bidirectionally primed pairs among 1,225 pairs of 50 TFs selected for interaction profiling.

#### Figure S2: Quality control metrics related to the dual-sgRNA Perturb-seq screen

A. Percentage of droplets called by cellranger or manual threshold as single cells assigned to 1, 2, or 3+ sgRNA identities in each GEM group. Median percentage assigned exactly 2 sgRNAs across GEM groups is annotated.
B. Gene expression quality control metrics for cells assigned exactly 2 sgRNA identities. Dashed lines indicate thresholds for filtering in final dataset used for analysis.
C. Distributions of Pearson correlation of transcriptional profiles (calculated in PCA space, 50 principal components) for pseudobulk populations assigned the same targets in different orientations (AB / BA), or for pairs of randomly chosen populations.
D. Number of cells maintained for analysis assigned to each sgRNA (316,630 total cells). Orientations are collapsed (cells assigned AB and BA are aggregated into one population). Colors denote double (AB) and single (A + NTC) perturbations. Leftmost point highlights NTC cells (NTC + NTC).

#### Figure S3: Comparison of double transcriptional profiles to RPE cell states observed *in vivo*

A. UMAP representations of Shao et al. 2026 RPE and choroid snRNA-seq atlas (left: 566,646 total cells, right: 69,244 RPE cells).
B. Distributions of state scores calculated from marker gene expression for double and single perturbations across five RPE states observed in atlas. Variance of state scores in doubles vs. singles compared via Levene's test and corrected via Benjamini-Hochberg: $*q < 0.05$, $**q < 0.01$.
C. Z-normalized expression of marker genes of *CST3*+, *INSYN2B*+, and *LGI1*+ RPE states delineated in atlas, in single and double perturbations involving the RPE identity factor *MYRF*.

#### Figure S4: AMD GWAS variants and heritability are enriched in RPE and RPE-1 cells

A. Enrichment vs. recall of 301 fine-mapped AMD risk variants (PIP > 0.1) by peak sets from different cell types in the RPE and choroid, using peak calls from scATAC-seq performed by Wang et al. Enrichment is relative to a background of 1000 Genomes common and low frequency variants (minor allele count ≥ 5). Dots show mean and error bars show 1.6 * standard error.
B. Heritability enrichment results computed using stratified LD score regression (S-LDSC) conditioned on the baseline model for the same peak sets in S6A. Heritability enrichment represents the proportion of heritability captured by SNPs overlapping the peak set divided by the proportion of total SNPs captured by the peak set. Bars show the enrichment and error bars show 1 standard error. $*p < 0.05$, $**p < 0.001$. *P*-values computed from standard errors derived from a block jackknife approach.
C. Heritability enrichment result from S-LDSC (same as S6B) showing the heritability enrichment for bulk ATAC-seq peaks called from our RPE-1 cell line. $*p < 0.05$. *P*-values computed from standard errors derived from a block jackknife approach.

**Figure S5: Further characterization of AMD risk loci that are revealed by perturbation-induced regions from Multiome Perturb-seq**

A. UpSet plot showing the variants revealed by scE2G peaks that are linked to genes, and the extent of their overlap with bulk or control RPE-1 peak sets. Peak sets defined as in Fig. 3B.
B. Histogram showing the number of perturbations that reveal the perturbation-specific credible sets ($n = 62$ credible sets total).

**Figure S6: Example of genetic epistasis hypothesis nominated by Multiome Perturb-seq**

Schematic showing how CRISPR activation of *MYRF* coordinately activates two downstream peaks that both overlap variants in distinct AMD credible sets. Coding both variants recessively, the AMD association of rs13278062 (PIP = 0.97) differed within the rs12061341 (PIP = 0.11) risk-genotype stratum, with the joint genotype associated with a lower proportion of AMD cases ($p = 1.5 \times 10^{-3}$, Wald test). Error bars show the 95% confidence interval based on a Wilson score interval.

**Figure S7: PoPS genes for AMD show preferential elevation of residuals compared to other traits.**

Comparison of the mean residual difference between PoPS genes and other genes across AMD and 113 other complex traits. Here, the difference between mean PCI score for PoPS genes and all other genes is computed across the 1, 5, 10, 15, 25, and $50^{th}$ percentiles of PoPS genes. Then, these differences are averaged to get the mean difference metric. X-axis is the traits ranked by mean difference, and Y-axis is the mean difference. CAD = coronary artery disease, T2D = type 2 diabetes, WBC = white blood cell count, TC = total cholesterol, and RBC = red blood cell count.

**Figure S8: Perturb-PoPS displays increased recall of Mendelian retinal disease (MRD) genes and learns convergence between polygenic AMD and MRD genes.**

A. Recall of MRD genes by different PoPS score variations. Perturb-PoPS - training features from our Perturb-seq data. PoPS-control - training features from control RPE-1 cells. PoPS - training features from original PoPS feature set. Recalls shown for either top 500 or top 1000 genes per score.
B. Recall of MRD genes by PoPS trained on singles and doubles features. Doubles - training features from singles and doubles. Singles - training features from only singles. Error bars - 1.6 * s.e. Recalls shown for either top 500 or top 1000 genes per score.
C. Enrichment and recall of MRD genes by Perturb-PoPS score trained with shuffled MAGMA scores. Recall and enrichment shown for top 500 genes prioritized by shuffled Perturb-PoPS. Error bars - 1.6 * s.e.

**Figure S9: Specification, performance, and attribution of gene-level interaction models**

A. Schematic of Mosaic architecture with descriptions of modules, inputs, and output.
B. Schematic of modeling approach for bilinear and simple nonlinear interaction models.
C. Performance (Pearson correlation between observed and predicted residuals for held-out genes in all pairs) of each modeling approach. Points show correlation within held-out genes in each model fold; bar represents mean across folds.
D. Relative performance (percent change from original fold's Pearson correlation on held-out genes) of Transformer-style model with peak features, single-TF expression, and single-TF ChromVAR scores ablated via replacement with training set mean in each fold. Bar represents mean across folds.

**Figure S10: Profiling of TF triples nominated by Mosaic**

A. Program attribution score for EGR motifs with respect to FOSB_SOX18 predictions vs. program residual in the EGR3 triple-TF condition. Programs are labeled with odds ratios and q-values of overlap with AMD PoPS genes as in Fig. 4D.

B. Program attribution score for sgIRF4 accessibility with respect to FOSB_SOX18 predictions vs. program residual in the IRF4 triple-TF condition. Programs labeled as above.
C. Distribution of residuals for 11,255 protein-coding genes detected in CREB3L3 single, FOSB_SOX18 double, and CREB3L3 triple conditions. Residuals for MRD genes *BEST1* and *CP* are highlighted with arrows.

### Supplemental Note: A regulatory state-dependent interpretation of the omnigenic model

The omnigenic model proposes that genetic effects on complex traits can be partitioned between relatively trait-proximal "core" genes and a much larger number of peripheral genes whose effects are mediated through regulatory networks[1,2]. In its formalized version, core genes are defined by relatively direct effects on phenotype, whereas peripheral genes can influence phenotype indirectly through *trans*-regulatory effects on core genes. The model further predicts that coordinated regulation of core genes can allow large numbers of individually weak regulatory effects to contribute substantially to complex-trait heritability.

Identifying core genes empirically is inherently difficult because directness of phenotypic effect is rarely known genome-wide. Phenotype-matched Mendelian disease genes provide one of the most widely used empirical proxies for candidate core genes, because large-effect mutations in these genes can produce disease phenotypes closely related to the corresponding complex disorder. Wray et al. (2018)[3], in discussing and critiquing the core-gene formulation of the omnigenic model, nevertheless highlighted genes underlying monogenic forms of disease as plausible examples of genes with direct phenotypic effects, while cautioning that common disease may involve a much broader and less sharply separable set of core genes than originally envisioned. Vuckovic et al.[4] subsequently provided an empirical implementation of this idea: they explicitly defined core genes as genes implicated in Mendelian blood disorders and showed that these genes were enriched among blood-trait GWAS signals, occupied central positions in co-expression networks, and were preferential targets of trans-eQTLs, properties consistent with predictions of the omnigenic model. For AMD specifically, Freund et al.[5] identified a substantial set of Mendelian retinal-disease genes matched to the AMD phenotype and demonstrated enrichment of these genes near common-variant AMD associations. We therefore refer to phenotype-matched Mendelian retinal-disease genes as "*candidate core genes*", rather than assuming that every such gene necessarily satisfies the mechanistic definition of a core gene.

Most earlier empirical tests of omnigenic predictions relied on steady-state regulatory relationships, particularly trans-eQTLs, trans-pQTLs, and coexpression networks. For blood-cell traits, Mendelian blood-disorder genes were found to occupy central positions in gene-regulatory networks[4] and to be preferential targets of *trans*-eQTLs and *trans*-pQTLs in whole blood[6,7]. Recently, Ota et al.[8] provided an experimental extension of this framework by combining gene-level loss-of-function burden effects with Perturb-seq-derived regulatory relationships. Their approach uses human genetic estimates of gene–trait effects together with experimentally measured gene-to-gene and gene-to-program regulation to reconstruct regulator–program–trait relationships, and further demonstrated that the utility of Perturb-seq for genetic interpretation depends strongly on studying a cellular context relevant to the trait. In this study, AMD provides a complementary setting in which to test whether these principles generalize to a non-hematopoietic, tissue-relevant regulatory system. RPE-1 provides an experimentally tractable retinal pigment epithelial context, while approximately 100 genes implicated in phenotypically matched Mendelian retinal disorders provide a substantial, independently defined candidate-core reference set.

Our study extends this Perturb-seq-based framing of the omnigenic model. Rather than using gene-level loss-of-function effects as the phenotypic anchor, we ask whether genome-wide common-variant AMD association signal converges on independently defined, phenotype-matched Mendelian retinal-disease genes through experimentally induced regulatory states. PoPS provides a natural framework for this question because it learns gene features that are systematically associated with polygenic GWAS signal rather than restricting inference to genome-wide significant loci[9]. By replacing conventional multi-tissue steady-state features with perturbation-derived gene features, Perturb-PoPS provides a genetic readout of which experimentally measured regulatory states most strongly expose the distributed genetic architecture of AMD. Mendelian retinal-disease genes are not used to train Perturb-PoPS and therefore provide an independent benchmark for testing whether perturbation-informed prioritization preferentially recovers candidate core genes.

Several complementary analyses in this work support this interpretation at different levels of the regulatory architecture. First, Pooled Combinatorial Interaction (PCI) measures each gene's aggregate sensitivity to non-additive regulation across TF combinations. Elevated PCI among Mendelian retinal-disease genes therefore indicates that candidate core genes are preferentially engaged by combinatorial regulatory interactions. Second, the simultaneous enrichment of Program 38 for genes prioritized by common-variant AMD genetics and for Mendelian retinal-disease genes provides program-level evidence for convergence between these independently defined genetic architectures within the same perturbation-responsive transcriptional state. Third, Perturb-PoPS shows that when genome-wide AMD association signal is interpreted through perturbation-derived gene features from individual TFs and combinatorial TF–TF perturbations, it preferentially prioritizes phenotype-matched Mendelian retinal-disease genes. Fourth, comparing Perturb-PoPS across perturbation classes tests whether the strength of polygenic-to-candidate-core convergence depends on regulatory state. Stronger recovery of Mendelian retinal-disease genes from combinatorial than single-perturbation features indicates that this convergence becomes more apparent in specific higher-order states. Fifth, at the level of individual genes, distinct TF pairs preferentially engaged different Mendelian retinal-disease genes, showing that state dependence affects not only the overall strength of candidate-core convergence but also the identity of the candidate core genes engaged in each regulatory state.

Taken together, these analyses do not establish that every Mendelian retinal-disease gene is a core gene, nor do they constitute a direct test of every prediction of the omnigenic model. Rather, they support a specific extension of its regulatory-network interpretation: the regulatory connections through which polygenic disease-associated biology converges on candidate core genes are conditional on cellular regulatory state. In this view, steady-state networks represent only one configuration of a broader regulatory landscape. Perturbation experiments can expose otherwise latent connections, while genome-wide polygenic information provides a means to determine which experimentally induced states are most relevant to disease.

### References


1. Boyle, E. A., Li, Y. I. & Pritchard, J. K. An expanded view of complex traits: from polygenic to omnigenic. *Cell* **169**, 1177–1186 (2017).
2. Liu, X., Li, Y. I. & Pritchard, J. K. Trans Effects on Gene Expression Can Drive Omnigenic Inheritance. *Cell* **177**, 1022-1034.e6 (2019).
3. Wray, N. R., Wijmenga, C., Sullivan, P. F., Yang, J. & Visscher, P. M. Common Disease Is More Complex Than Implied by the Core Gene Omnigenic Model. *Cell* **173**, 1573–1580 (2018).
4. Vuckovic, D., Bao, E. L., Akbari, P., Lareau, C. A., Mousas, A., Jiang, T., Chen, M.-H., Raffield, L. M., Tardaguila, M., Huffman, J. E., Ritchie, S. C., Megy, K., Ponstingl, H., Penkett, C. J., Albers, P. K., Wigdor, E. M., Sakaue, S., Moscati, A., Manansala, R., Lo, K. S., Qian, H., Akiyama, M., Bartz, T. M., Ben-Shlomo, Y., Beswick, A., Bork-Jensen, J., Bottinger, E. P., Brody, J. A., van Rooij, F. J. A., Chitrala, K. N., Wilson, P. W. F., Choquet, H., Danesh, J., Di Angelantonio, E., Dimou, N., Ding, J., Elliott, P., Esko, T., Evans, M. K., Felix, S. B., Floyd, J. S., Broer, L., Grarup, N., Guo, M. H., Guo, Q., Greinacher, A., Haessler, J., Hansen, T., Howson, J. M. M., Huang, W., Jorgenson, E., Kacprowski, T., Kähönen, M., Kamatani, Y., Kanai, M., Karthikeyan, S., Koskeridis, F., Lange, L. A., Lehtimäki, T., Linneberg, A., Liu, Y., Lyytikäinen, L.-P., Manichaikul, A., Matsuda, K., Mohlke, K. L., Mononen, N., Murakami, Y., Nadkarni, G. N., Nikus, K., Pankratz, N., Pedersen, O., Preuss, M., Psaty, B. M., Raitakari, O. T., Rich, S. S., Rodriguez, B. A. T., Rosen, J. D., Rotter, J. I., Schubert, P., Spracklen, C. N., Surendran, P., Tang, H., Tardif, J.-C., Ghanbari, M., Völker, U., Völzke, H., Watkins, N. A., Weiss, S., VA Million Veteran Program, Cai, N., Kundu, K., Watt, S. B., Walter, K., Zonderman, A. B., Cho, K., Li, Y., Loos, R. J. F., Knight, J. C., Georges, M., Stegle, O., Evangelou, E., Okada, Y., Roberts, D. J., Inouye, M., Johnson, A. D., Auer, P. L., Astle, W. J., Reiner, A. P.,

Butterworth, A. S., Ouwehand, W. H., Lettre, G., Sankaran, V. G. & Soranzo, N. The Polygenic and Monogenic Basis of Blood Traits and Diseases. *Cell* **182**, 1214-1231.e11 (2020).
